# A synthetic protein-level neural network in mammalian cells

**DOI:** 10.1101/2022.07.10.499405

**Authors:** Zibo Chen, James M. Linton, Ronghui Zhu, Michael B. Elowitz

## Abstract

Artificial neural networks provide a powerful paradigm for information processing that has transformed diverse fields. Within living cells, genetically encoded synthetic molecular networks could, in principle, harness principles of neural computation to classify molecular signals. Here, we combine de novo designed protein heterodimers and engineered viral proteases to implement a synthetic protein circuit that performs winner-take-all neural network computation. This “perceptein” circuit includes modules that compute weighted sums of input protein concentrations through reversible binding interactions, and allow for self-activation and mutual inhibition of protein components using irreversible proteolytic cleavage reactions. Altogether, these interactions comprise a network of 310 chemical reactions stemming from 8 expressed protein species. The complete system achieves signal classification with tunable decision boundaries in mammalian cells. These results demonstrate how engineered protein-based networks can enable programmable signal classification in living cells.

**One-Sentence Summary:** A synthetic protein circuit that performs winner-take-all neural network computation in mammalian cells

## Introduction

Cells are classification machines. Using circuits of interacting genes and proteins, they make qualitatively distinct decisions in response to the levels or dynamics of multiple input signals. For example, p53 functions as a tumor suppressor by classifying the types and levels of stress the cell encounters, and inducing senescence or cell death in response (*1*). In the context of development, cells in the neural tube take on specific progenitor fates by classifying the levels of BMP and Hedgehog signaling (*2*). In the human immune system, the classification of multiple cytokine inputs can control the fate of a T cell (*3*). The ability to program synthetic signal classification systems could facilitate engineered gene and cell therapies by allowing cells to robustly distinguish diseased and normal cell states (*4*). For this reason, a major goal of synthetic biology has been to design synthetic classification circuits that could function in living cells, respond to different types of signals as inputs and control cellular functions as outputs (*5*).

One of the most powerful circuit architectures for classification is the winner-take-all neural network (*6*). In these systems, an output neuron is ON if and only if the weighted sum of its inputs exceeds that of all other neurons in the output layer (Fig. 1A, B). This architecture has several benefits. First, it offers a compact mechanism for signal classification, requiring only a single layer neural network. Second, it ensures that outputs are all-or-none (Fig. 1B). Finally, it allows one to alter the decision boundary simply by tuning weights (Fig. 1C).

**Figure 1.**
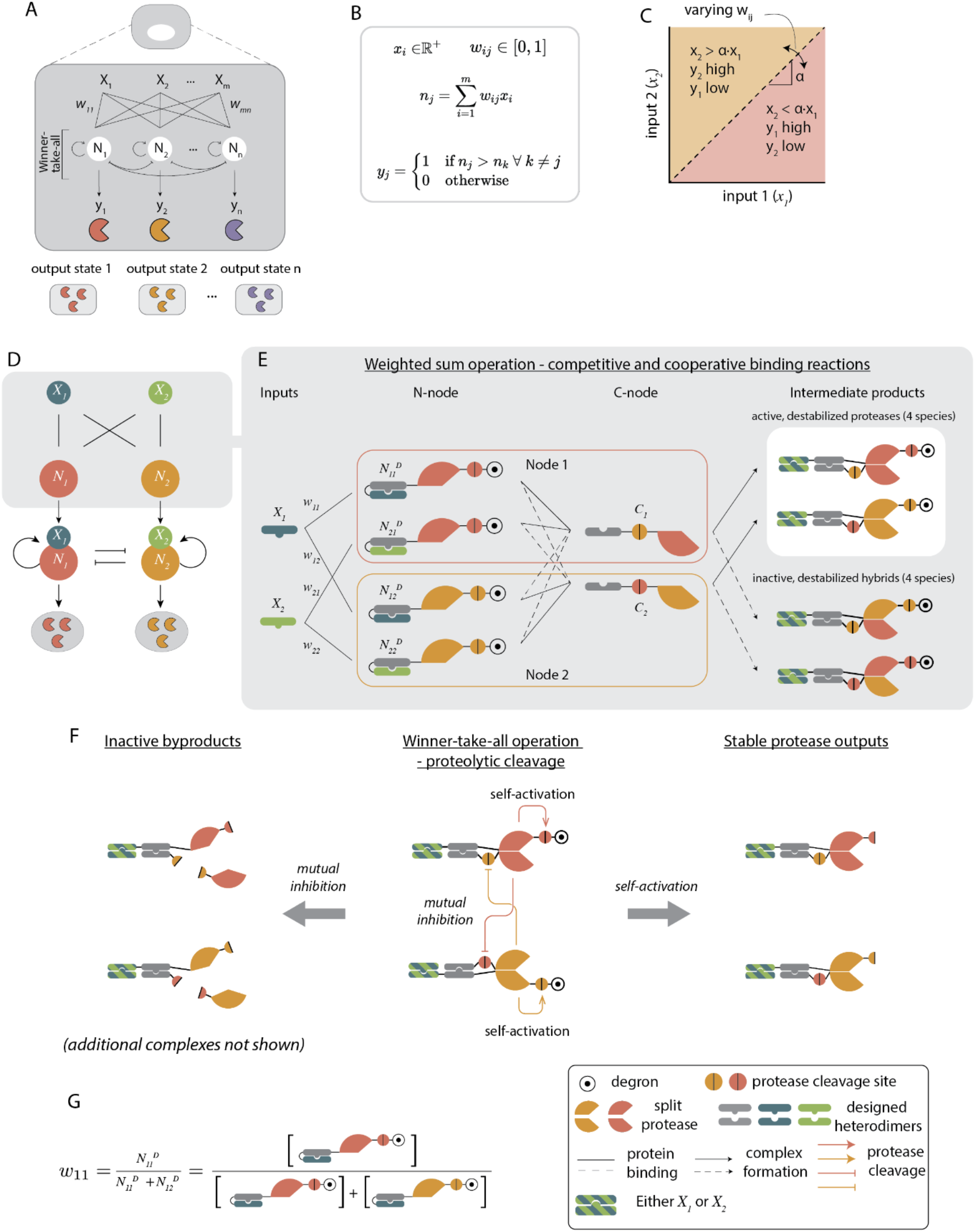
Winner-take-all neural network computation can be implemented using engineered proteins. (A) A winner-take-all neural network, operating inside cells, would use a set of interacting proteins (N_j_) to activate exactly one of its outputs (colored proteases, y_j_) depending on the relative values of its inputs (X_i_). (B) Formal description of the system in (A). The network consists of m inputs (X_i_), each taking positive real values. The inputs interact with the n nodes (N_j_) with weights w_ij_ (1 i m, 1 j n, 0 w_ij_ 1) connecting input X_i_ to neuron N_j_. Each neuron performs weighted sum operations to integrate the input signals it receives, and winner-take-all is achieved through self-activation and mutual inhibition. The output y_j_ from a neuron N_j_ is active only if its weighted sum is greater than that from any other neuron. (C) The decision boundary, a, for a 2-input, 2-neuron network can be tuned by varying the weights w_ij_. (D) In a 2-input, 2-node circuit, each input protein activates either node protein by forming input-node complexes. Such complexes then undergo self-activation and mutual inhibition to perform the winner-take-all computation. The final state of the system is defined by the abundance of the active node. (E) The weighted sum operation is carried out through competitive and cooperative binding. The two inputs are de novo designed orthogonal DHDs (X_1_ and X_2_). Each node consists of two groups of proteins: the N-nodes, where the cognate binding partners of X_1_ and X_2_ are caged by a genetically fused DHD caging domain, and further linked to the N-terminal half of a protease, its cleavage site, and a DHFR degron; the C-nodes, made from the cognate binding partner of the DHD caging domain in primary half-nodes, fused to the cleavage sequence of the other protease, and the C-terminal half of a protease. The inputs, N-node, and C-node bind cooperatively, such that neither of the two proteins can bind with high affinity without the third protein. They also interact competitively, such that the N-nodes compete to bind to the input protein, and the C-nodes. Two types of intermediate products result from these 3-way binding events: the active but destabilized proteases where the two protease halves reconstitute a functional protease, or the inactive and destabilized hybrids where the two protease halves do not match. The blue and green stripes indicate that the DHD domains can be either blue (X_1_) or green (X_2_). (F) Winner-take-all operation is achieved through two types of reactions: mutual inhibition, where each protease can inactivate the opposite protease type by cleaving its C-terminal half proteases off the C-nodes; self-activation, where the intermediate DHD-protease complexes cleave off DHFR degrons from their N-nodes, converting them to stable proteases, which can in turn activate and inhibit other protease complexes. (G) The weights connecting inputs to neurons are set based on the abundance of each primary half-neuron complex. For example, w_11_, the weight that connects input X_1_ to Neuron 1, is defined as the concentration of the N_11_ N-node divided by the sum of the concentrations of all N-nodes that can potentially bind to X_1_, in this case N_11_ + N_12_.

Previous theoretical work suggested specific schemes for engineering biochemical neural computation systems (*7*-*11*). Experimentally, efforts to build synthetic classification systems have resulted in DNA classifiers in test tubes (*12, 13*), and miRNA-based classifiers in mammalian cells (*14, 15*). In contrast, a protein level classifier would offer several advantages: it could be expressed transiently in cells, interface directly with endogenous inputs and outputs, trigger different output pathways depending on input state, and should in principle work in diverse cell types without relying on endogenous transcriptional regulations. More generally, a protein-based neural network would also test the ability to construct sophisticated computational devices out of interacting proteins (*16*).

A key challenge in designing a protein-based neural network has been the increased difficulty of programming specific binding interactions and conditional activity using proteins compared to nucleic acids. Recently, however, sets of modular protein interaction domains have been developed, including de novo designed heterodimers (DHDs) (*17*-*20*). Additionally, multiple groups have used conditional reconstitution of split viral proteases as a way to control protein activities (*21*-*24*). Here, taking advantage of these advances, we designed a protein based winner-take-all comparator circuit using DHDs and split proteases that senses and processes input signals. This circuit accurately compared the relative levels of two inputs in living mammalian cells, with a tunable decision boundary. This synthetic protein neural network provides a foundation for rationally engineering protein classification in living cells.

## Results

### System design

Winner-take-all dynamics can be achieved with molecular components that exhibit three key features (Fig. 1D): First, they should respond to input molecules with tunable strengths, analogous to weights in neural networks, to enable weighted summation of inputs. Second, they should be capable of mutual inhibition, allowing the elimination of less abundant species. Third, they should be able to self-activate, so that surviving species can amplify their own activity (*25*).

We created fusion proteins that possess these features by genetically combining de novo designed protein heterodimers (DHDs) (*17*), domains from viral proteases such as Tobacco Etch Virus (TEV) and Tobacco Vein Mottling Virus (TVMV) (Fig. 1E), and the dihydrofolate reductase (DHFR) degron. Briefly, the proteases are split into two inactive domains, whose reconstitution into an active protease can be controlled by attached DHD domains (*21, 22*). The intermolecular dimerization of these DHD domains can be regulated by input DHDs that disrupt a competing inhibitory self-caging interaction (Fig. 1E) (*26*). The strength of this input binding interaction, analogous to a weight in a neural network, is tunable by varying the concentrations of DHD-protease fusions (Fig. 1G). Once reconstituted, proteases can inactivate one another by cleaving off attached dimerization domains, achieving mutual inhibition (Fig. 1F). Finally, the same proteases can also self-activate by cleaving off degrons that would otherwise destabilize them (Fig. 1F). Together, these components generate the key interactions required for constructing a set of protein-based winner-take-all neural networks we term *perceptein networks*, in loose analogy with the perceptron, a foundational artificial neural network architecture (*27*).

In the language of neural networks, the perceptein network comprises a layer of inputs (DHD input proteins, X_i_), as well as a layer of nodes (N_i_) that process information from inputs and couple to outputs (Fig. 2). In this molecular implementation, each node consists of two parts, labeled N-nodes and C-nodes. The N-nodes are composed of N_jk_^D^ or N_jk_, consisting of N-terminal protease halves fused to DHDs, with (N_jk_^D^) or without (N_jk_) an attached degradation tag. The C-nodes, labeled C_l_, contain C-terminal protease halves fused to DHDs (Fig. 1E). In the simple case of two inputs and two outputs, all indices range from 1 to 2, such that the whole system consists of two X_i_, four N_jk_^D^, and two C components, or eight proteins altogether. Throughout the text, N_i_ refers to the node consisting of N-node and C-node proteins, while N_jk_^D^ and N_jk_ refers to individual N-node proteins.

**Figure 2.**
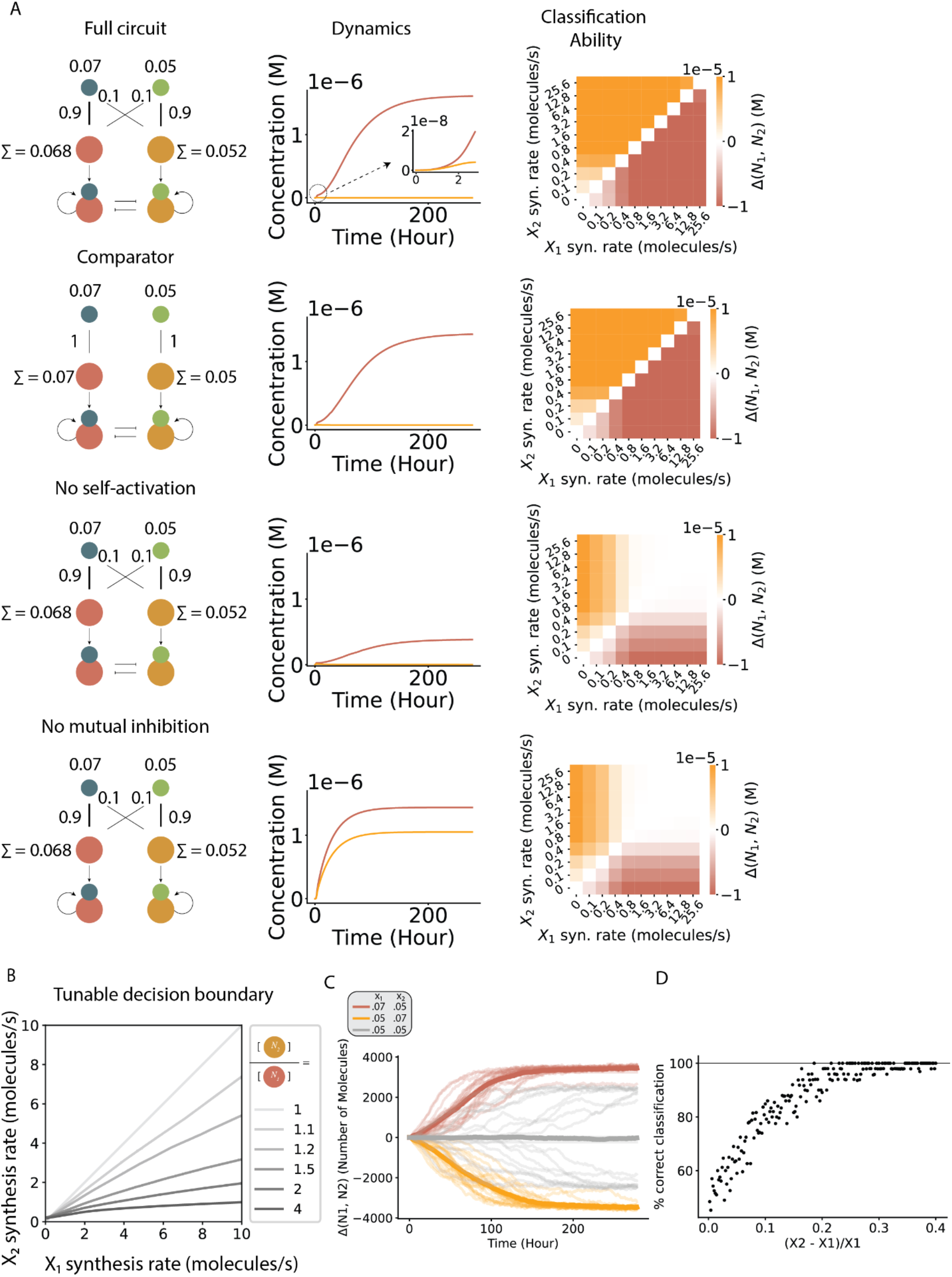
Simulated two-input circuits perform winner-take-all classification. (A) Simulations of circuit variants (left) reveal circuit dynamics (middle) and classification ability (right). Input values, weights, and weighted sums (denoted L) are indicated on the circuit diagrams, with larger weights represented by thicker lines. Each cell in the heatmap represents the difference between active N_1_ and N_2_ proteases at steady state. Both the full circuit and the comparator are able to classify across the full range of input levels, while circuits lacking self-activation or mutual inhibition only classify within a limited input range. (B) The decision boundary (gray lines) of the comparator circuit can be tuned by varying the relative levels of the two node proteins, N_11_^D^ and N_22_^D^. (C) Stochastic simulations of the comparator. Twenty simulations were performed for each condition (light traces), and their average traces are plotted in dark lines. Colors indicate the input levels (legend). See supplementary materials for simulation methods. (D) Percentage of 50 equivalent simulations that correctly classify inputs as a function of the concentration difference between the two inputs. Input X_1_ is fixed at 0.05 molecules/s. Even with stochasticity, input differences of at least 20% classify correctly ∼95% of the time.

Our design enables weighted summation of inputs. The i^th^ input species, X_i_, is partitioned among different potential binding partners N_jk_^D^ (for which i = j) in proportion to their relative abundances. The effective weights, w_ij_, are thus determined by the abundances of the corresponding N_jk_^D^ components. For example, in the case of 2 inputs and 2 outputs, w_11_ is defined as the concentration of N_jk_^D^ divided by the sum of the concentrations of N_jk_^D^ and N_jk_^D^ (Fig. 1G). Binding to an input uncages (*26*) the DHD domain fused to N_jk_^D^ components, exposing the N_jk_^D^ binding domain (gray), which can interact with C_l_ domains to produce either a functional protease (if k = l) or a non-functional hybrid of mismatched protease halves (if k ≠ l) (Fig. 1E). Each input can, in this way, generate both functional reconstituted proteases as well as non-functional hybrids. Input summation occurs because the total set of functional reconstituted proteases of a given type in general includes contributions from all inputs.

The design also enables self-activation and mutual inhibition. To achieve self-activation, functional proteases can further activate their own activity by cleaving off a degron that is present on the N_jk_^D^ components of the same protease type, which would otherwise cause rapid protease degradation (Fig. 1F). The functional reconstituted proteases of each type can also inactivate other functional proteases by cleaving off DHDs on their C_l_ domains (Fig. 1F).

### Modeling and simulation

To understand whether this design could produce winner-take-all behavior within physiologically relevant parameter regimes, we first simulated its response to a matrix of input values (X_i_ concentrations) (Supplementary Text). Briefly, inputs X_i_ bind cooperatively to N_jk_^D^ and C_l_ to form trimeric complexes that reconstitute either functional proteases or nonfunctional hybrids with mismatched protease halves. To model this cooperative binding process, we divided the process of trimer formation into two steps. First, X_i_ binds to N_ij_^D^ to form an unstable dimer with fast off rates. This step also exposes the binding site for C_l_. In the second step, this dimer binds to C_l_ to form stable trimeric complexes with slower off rates (*26*). Complexes with matching protease halves are assumed to reconstitute active protease. Each reconstituted protease can cleave its various cognate target sites on other protein components. All the while, all protein building blocks are continuously synthesized at constant rates and degraded at different rates depending on whether they contain a degron. Finally, we estimated physiologically reasonable values for protein synthesis and degradation rates, protease catalytic rates, and other biochemical parameters using references in the BioNumbers (*28*) database (Table 1).

With these parameter values, the simulated circuit exhibited the desired winner-take-all classification behavior (Fig. 2A, first row). When X_1_ exceeded X_2_, the N_1_ protease activation approached its maximum possible level over timescales of ∼100h. However, the decision appeared to be made much earlier, as large fold differences between N_1_ and N_2_ were apparent within 3 hours (Fig. 2A, first row inset). Simulations further revealed that inner-take-all classification functions across a broad range of absolute concentrations for X_1_ and X_2_, and remained accurate even for differences as small as 10% between the two ligand concentrations (Fig. S1C).

To understand which features of the circuit are necessary or sufficient for classification, we also analyzed circuit variants lacking cross-activation by inputs (Fig. 2A, second row), self-activation (Fig. 2A, third row), or mutual inhibition (Fig. 2A, fourth row). Removing cross interactions (i.e., removing N_12_^D^ and N_21_^D^) produced a simplified circuit design we term the comparator (Fig. 2A, second row) that is still capable of winner-take-all classification. Removing self-activation retains all-or-none behavior at lower input levels, but at a lower dynamic range of outputs (Fig. 2A, third row). At higher input levels, it loses classification ability altogether. By contrast, removing mutual repression accelerated the response of the circuit but eliminated the all-or-none output behavior, and also lost classification ability at higher input levels (Fig. 2A, bottom row). Therefore, both self-activation and mutual inhibition are indispensable for winner-take-all computation in this architecture.

We next analyzed sensitivity to parameter values within the circuit. Protease catalytic rates and protein degradation rates both had strong effects on classification accuracy (Fig. S1A, B). Lower values of k_cat_ led to reduced cleavage, hindering self-activation and cross-inhibition and therefore decreasing classification performance. However, as long as k_cat_ values for both proteases were above 0.04 sec^-1^, the circuit operated correctly, identifying the larger of the two inputs (Fig. S1A). Additionally, the winner-take-all behavior was more pronounced at higher degradation rates, corresponding to stronger degrons in the experimental circuit (Fig. S1B, x-axis). By suppressing background protease activity, degradation amplifies the effects of self-activation.

In general, different inputs can vary over different ranges. An ideal circuit would allow one to tune the decision boundary to match the scales of the inputs. In these circuits, the decision boundary can in fact be tuned by varying weights, represented here as component concentrations. In the comparator, when N_11_^D^ and N_22_^D^ are fixed at the same level, the circuit compares input levels without bias. On the other hand, varying the relative abundances of N_11_^D^ and N_22_^D^ allows the construction of biased comparators, where N_2_ is ON only when X_2_ > X_1_ ·α, where α depends on the relative levels of N_11_^D^ and N_22_^D^. Simulations confirmed that the circuit could implement such tunable decision boundaries (Fig. 2B).

Within cells, stochastic fluctuations can strongly impact circuit behaviors (*29*). Stochastic simulations of the circuit revealed that both the full network and the comparator circuits could function accurately despite such noise (Fig. 2C, Fig. S1D-F). Even with a difference in input values of only 40% (e.g. X_1_=0.05, X_2_=0.07), no “reversal” events were observed, as shown by individual traces (red and orange lines, Fig. 2C). On the other hand, when starting from equal inputs (X_1_=X_2_=0.05), individual trajectories converged over a slower timescale to either of the two output states, with an average response of neither (dark gray line, Fig. 2C), showing that the output is bistable. To analyze the sensitivity to input differences in the presence of noise, we varied X_1_-X_2_ and analyzed the fraction of correct decisions. This analysis suggested that differences of only 20% were sufficient for accurate classification more than 95% of the time (Fig. 2D).

These results suggest that the perceptein architecture should function across a broad, biologically plausible range of parameter values. On the other hand, the perceptein produces an enormous number of distinct molecular species. Even in the smallest implementation, involving two inputs and two neurons (Fig. 2A, first row), starting from just 8 protein species, the system generates (by cleavage and protein complex formation) 158 unique proteins and protein complexes, that participate in 310 distinct chemical reactions, including protein binding, synthesis, degradation, and protease cleavage (Supplementary Text). Modeling these reactions required a set of 158 ordinary differential equations containing 1238 terms. This complexity, which exceeds that of most previous synthetic biological circuits, provokes the question of whether the system could actually function in mammalian cells.

### Experimental validation

We constructed the set of 6 perceptein components and 2 input proteins necessary to implement the full circuit (Fig. 3A, S2). We chose the split tobacco etch virus protease (TEVP) and tobacco vein mottling virus protease (TVMVP) (*21*) as the two orthogonal proteases, and genetically fused the split halves to DHD domains, protease cleavage sequences, and degradation domains for controlled reconstitution of full proteases (Fig. 3A, S2, Table 2). In order to test many combinations of components and expression levels by co-transfection, each protein was encoded on a distinct plasmid. To measure the output of the circuit, we engineered a stable HEK293 reporter cell line containing a multi-cistronic construct co-expressing two fluorescent proteins-mCitrine and mCherry-each tagged with a cleavage-activated N-degron for either of the two input proteases (*21*) (Figure 3B). We verified that each protease variant exclusively reduced fluorescence from its target reporter (Figure S3A). Together, these constructs and the reporter cell line permitted rapid, iterative testing of circuit designs.

**Figure 3.**
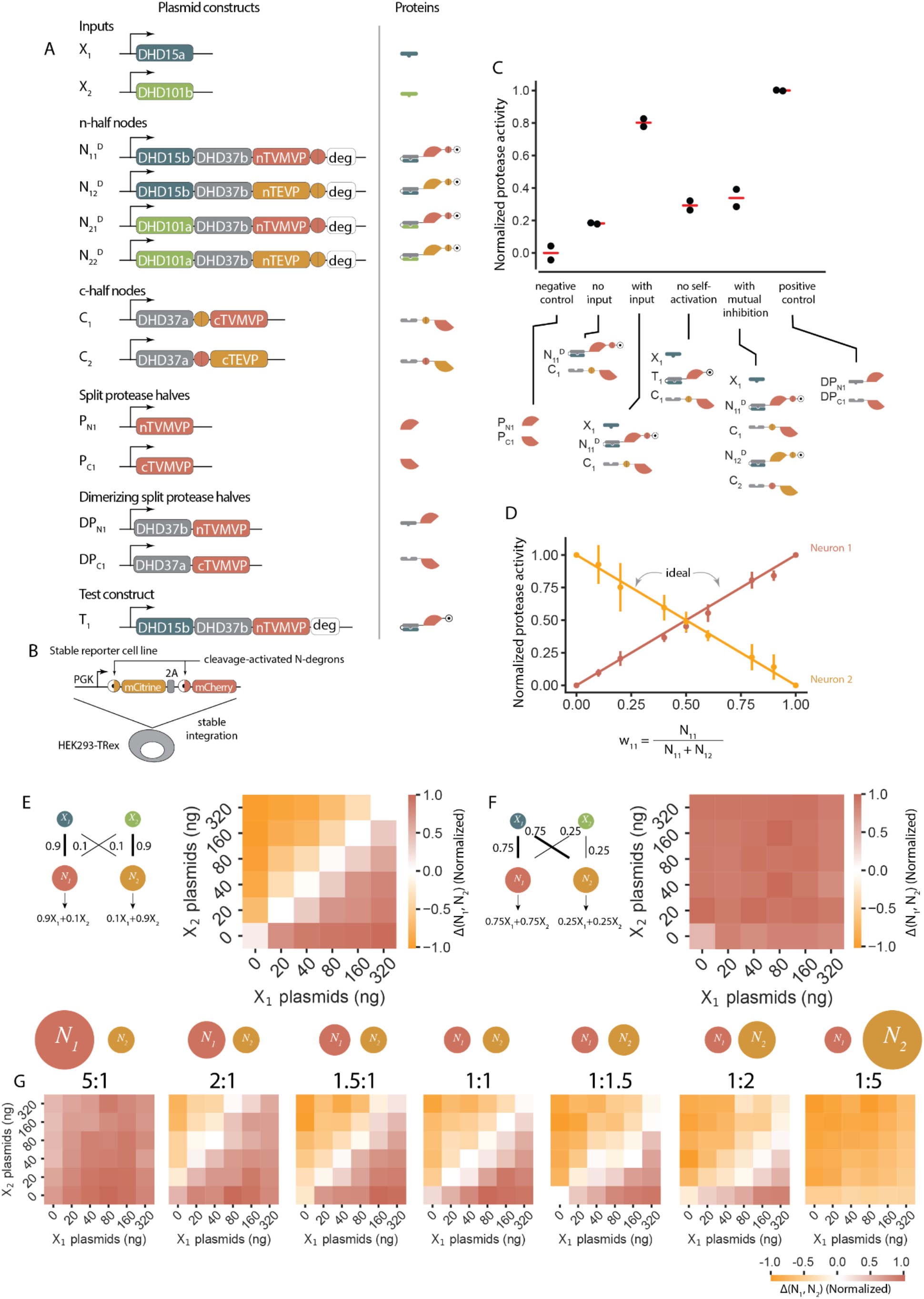
The winner-take-all neural network circuit classifies inputs in mammalian cells. (A) Plasmids and the encoded protein constructs. All plasmids use the human cytomegalovirus promoter (PCMV) to express each engineered protein. Deg denotes DHFR degron. Orange and yellow circles denote cleavage sites for corresponding proteases. DHDs are designed heterodimerizing proteins. A schematic representation of each fusion protein is shown to the right of the construct. (B) The stable reporter cell line constitutively co-expresses mCitrine and mCherry fluorescent proteins that can be cleaved at the N-terminus by TEVP and TVMVP, respectively, to reveal N-terminal degrons that destabilize the fluorescent proteins. PGK, 3-phosphoglycerate kinase promoter. (C) Engineered protease can respond to inputs, self-activate, and mutually inhibit. Normalized protease activities under different experimental setups indicate expected functions. (D) Testing the weight multiplication module by fixing input X_1_ and varying nodes N_11_^D^ and N_12_^D^. Ideal behaviors are shown in solid lines, experimental data points are mean ± s.d from three biological repeats. (E) A fully connected 2-input, 2-neuron circuit that compares relative input levels (left). (F) A fully connected 2-input, 2-neuron circuit that, by construction, should always result in N_1_ being the winner. (G) The decision boundaries of a two-input comparator can be tuned by varying the ratios of N_1_ to N_2_ protein concentrations. Data in E-G are averages of two biological replicates.

Using these components, we experimentally validated each module of the winner-take-all neural network, starting from the top of Figure 1D. Each experiment was performed with varying amounts of input plasmids to identify combinations that maximize dynamic range and minimize background activities. First, we asked whether inputs could trigger reconstitution of corresponding protease activities, as depicted in Figure 1E. Co-transfecting input X_1_ with cognate N-and C-node proteins inactivated the corresponding fluorescent protein reporter, consistent with reconstitution of the corresponding protease (Figure 3C, S3C, S3D). Input-triggered protease activities were comparable to positive controls, consisting of split protease halves fused to heterodimerizing domains. In the absence of input, reporter levels were similar to those in a negative control consisting of split protease halves lacking DHD domains. These results suggest that inputs can reconstitute cognate protease activities.

Next, we focused on the weight multiplication step, which should ideally distribute input proteins based on the relative abundance of N-node proteins (Fig. 1G). In order to obtain a homogeneous distribution of constructs in each cell, we used mRNAs encoding test constructs for transfection. We transfected cells with varying ratios of N_11_ and N_12_, while keeping X_1_ constant. The amount of activated (protease reconstituted) N_1_ and N_2_, which determines the fluorescence of mCitrine and mCherry, should be linearly dependent on the ratio of N_11_ and N_12_ expression levels. Flow cytometry analysis confirmed that the changes in fluorescence followed a linear trend consistent with the concentration ratios of transfected N_11_ and N_12_ constructs (Fig. 3D).

Once activated, the perceptein components can self-activate and mutually inhibit (Fig. 1F). Self-activation involves protease cleavage-dependent removal of a fused DHFR degron from the half protease, stabilizing proteases of its own kind. To evaluate the extent of self-activation, we transfected HEK293 cells with plasmids encoding either N-node constructs with a protease cleavable degron, or similar negative control constructs lacking the cleavage site that are therefore unable to self activate. Flow cytometry revealed a 5 fold change in protease activity upon self-activation. By contrast, the negative control remained close to the background (Fig. 3C and S3E). Mutual inhibition interactions (Fig. 1F) also functioned as expected. As shown in Figure 3C, protease activities of the N_1_ node were strongly repressed by an excess of the N_2_ node components (N_12_^D^ and C_2_). Mutual inhibition worked similarly in the opposite direction when there was more N_1_ than N_2_ (Fig. S3E). Taken together, these results indicate that each module of the full circuit can function individually.

To test whether the full circuit behaved as predicted, we co-transfected reporter cells (Fig. 3B) with varying concentrations of the inputs, fixed amounts of the N-and C-half-node proteins (in a multicistronic manner), and a BFP co-transfection marker (Figure S2), and read out reporter fluorescence. We set the concentrations of the N_11_^D^ and N_22_^D^ plasmids 9 times higher than the concentrations of the N_12_^D^ and N_21_^D^ plasmids to put the circuit into a comparative regime (Fig. 3E, left). We normalized protease activities based on fluorescence (Materials and methods), and plotted the differences in normalized protease activities in N_1_ and N_2_. As expected, the output was positive when X_1_ exceeded X_2_ and negative when X_2_ exceeded X_1_, with minimal response at equal input concentrations (Fig. 3E, right). The output became more binary with greater absolute difference between the two inputs, approaching the binary response observed in simulations. Additionally, varying the relative levels of perceptein components resulted in a biased comparator where, in agreement with prediction (Fig. 3F, left), Node 1 was the winner regardless of the two input values (Fig. 3F, right). These results show that the full circuit can compare the relative levels of two inputs.

In the model, eliminating the cross interactions, X_1_-N_2_ and X_2_-N_1_ altogether, leads to a simpler comparator regime that should allow input classification with a decision boundary whose position can be tuned by modulating the relative expression levels of the perceptein components (Fig. 2A, second row). To test this capability, we transfected cells with varying relative levels of the N_11_^D^ and N_22_^D^ plasmids, while omitting the N_11_^D^ and N_21_^D^ plasmids entirely. At a 1:1 ratio of N_11_^D^:N_22_^D^, we observed similar classification behavior to that seen with the full circuit (Fig. 3E). Modulating the relative levels of N_11_^D^ and N_22_^D^ shifted the decision boundary, similar to predictions (Fig. 3G). These results support the ability of the circuit to enable tunable classification.

### The perceptein can produce scalable classification

How well can this protein-based neural network scale? To address this question, we simulated higher dimension comparators, each composed of *m* inputs and *m* nodes (Fig. 4A). For each value of *m*, we simulated the response to a matrix of input values. Larger systems retained classification ability, despite some loss of output dynamic range (Fig. 4B, C). Overall circuit complexity, measured by the number of chemical reactions, scaled approximately linearly with the size of the comparator, *m* (Fig. 4C).

**Figure 4.**
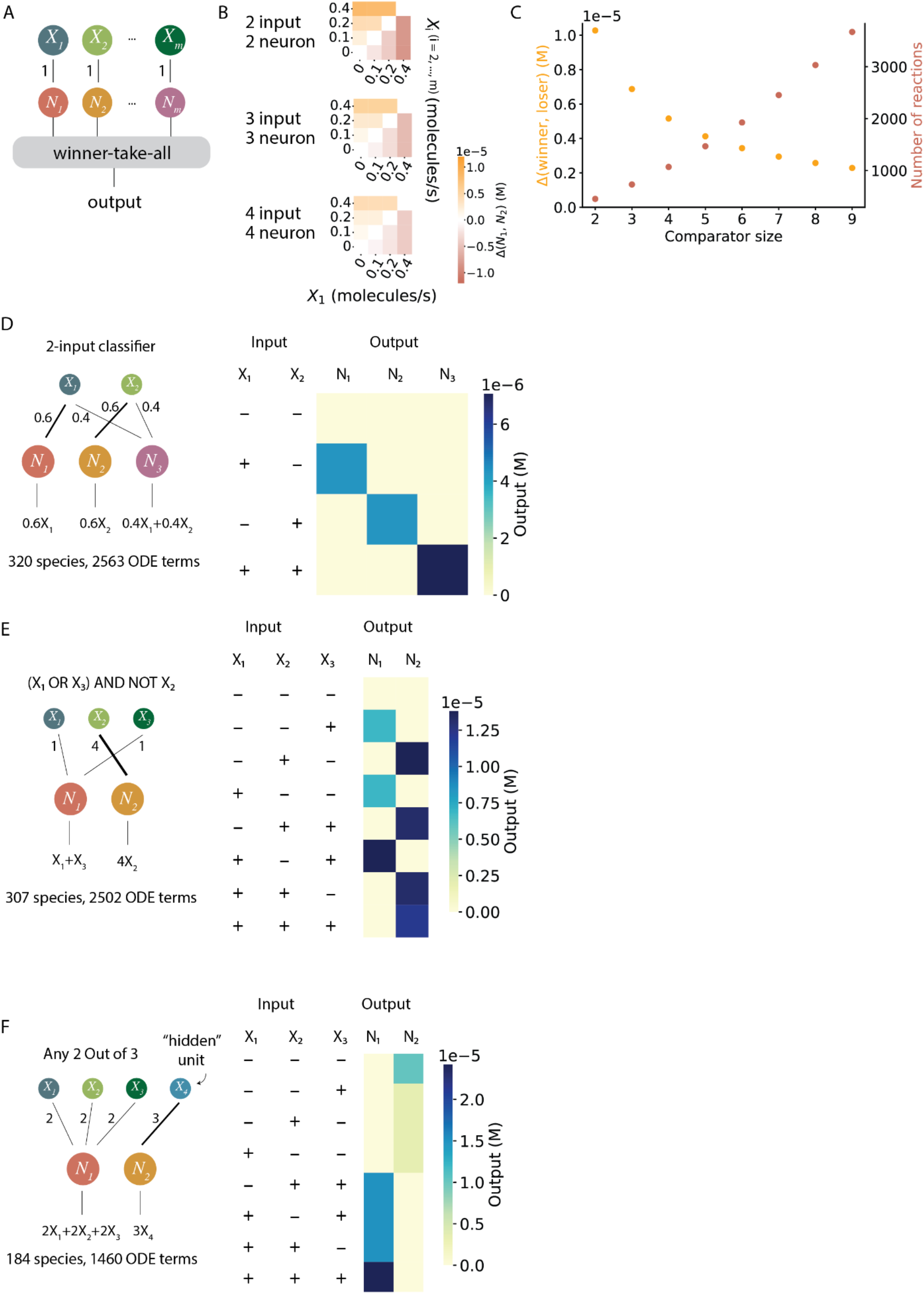
Scaling of the winner-take-all circuit. (A) An m-input, m-neuron comparator circuit. (B) Classification abilities of comparators that take 2, 3, and 4 inputs. As the size of the comparator increases, the ability to compare relative input levels is retained, while dynamic range is reduced due to the increased total number of substrates for each protease. (C) The number of reactions in a comparator circuit increases roughly linearly with its size, as the circuit dynamic range decreases. (D) A 2-input classifier can generate distinct responses to all 4 input states. (E) A 3-input winner-take-all circuit performs the (X_1_ OR X_3_) AND NOT X_2_ calculation. Neuron 1 wins if the condition is met. (F) A 3-input winner-take-all circuit performs “Any 2 out of 3” logic. A fourth “hidden unit” input was added to set the threshold and make the circuit more compact. Neuron 1 wins if the condition is met.

Finally, simulations showed that the perceptein system can also perform more complex types of classification. For example, by adding a third node to the two-input classifier, one can obtain a winner-take-all response in which nodes 1 and 2 respond to X_1_ or X_2_ alone, while node 3 responds only to the presence of both (Fig. 4D). Conversely, with three inputs and two nodes one can, in a single layer system, compute composite functions such as (X_1_ OR X_3_) AND NOT X_2_ that require multiple layers of conventional Boolean logic gates (Fig. 4E). One can also augment the system with “hidden” units that establish thresholds for the true inputs in order to compute even more complex functions. For example, by adding a single hidden unit to the 3-input, 2-node system, one can obtain a circuit that computes an ANY 2 OUT OF 3 function, where node 1 is activated when at least two of the three inputs are present (Fig. 4F). Together, these results show that the perceptein system can in principle be scaled and extended to solve a broader variety of classification problems.

## Discussion

Here, inspired by the classic winner-take-all neural network design, we introduced the perceptein architecture for protein level winner-take-all classification within living mammalian cells. The circuit design is based on three design principles: First, three-way cooperative binding interactions enable input-dependent protease activation (*26*), producing the species on the right in Fig. 1E. These cooperative interactions are in turn enabled by the use of fusions of de novo designed heterodimers. Analogous functions could in principle be achieved with naturally cooperative protein binding systems such as those in the N-WASP system (*30*). However, the de novo designed proteins offer a larger repertoire of distinct binding specificities, and minimize unintended interactions with endogenous cellular proteins. Second, the design takes advantage of the irreversibility of protease-based cleavage, in combination with degrons (*21*). Self-activation involves proteolytic degron removal, while mutual inhibition is achieved by proteolytic inactivation (Fig. 1F). We note that other mechanisms, such as post translational modification, could also be exploited to allow orthogonal signal processing and enable multi-layer protein networks. Third, perceptein components further exploit molecular competition between protease halves. Splitting proteases and letting them compete to form productive or inactive complexes achieves two functional goals: it makes each protease activity dependent on a cognate input, and it supports the winner-take-all behavior, as protease halves effectively quench the activity of their non-matching partners (Fig. 1E). Combining these principles successfully enabled two-input winner-take-all classification in mammalian cells (Fig. 3).

A key feature of neural computation is the ability to modulate input-output functions by tuning weights in an existing network (*27*). Analogously, varying the expression levels of perceptein circuit components such as N_11_^D^ and N_12_^D^ systematically shifted the decision boundary in the classification circuit without requiring additional protein components (Fig. 2B, 3G). Because tuning can be achieved through expression, this design allows cells to control their own decision thresholds by modulating gene expression.

The perceptein architecture can be scaled up to perform higher dimensional classification tasks. A fully connected network (Fig. 2A, first row) with p inputs and q neurons grows multiplicatively, requiring a total of p + q + pq distinct proteins. However, in the case of the simpler comparator architecture (Fig. 2A, second row), elimination of cross interactions between inputs and node proteins leads to linear scaling, with p + q starting components. When considering scaling, it is interesting to also consider the remarkably large number of different protein species and complexes (e.g. 158 for the 2-input, 2-output system) that are generated in the operation of this system. Effectively, this large number of molecular species is “compressed” into the much smaller number (e.g. 8 for the 2-input, 2-output system) of starting protein species from which they are generated. This is reminiscent of the way certain protein families can produce huge diversities of protein products through alternative splicing (*31*), proteolytic cleavage of pro-proteins (*32*), or combinatorial assembly of alternative multimeric complexes (*33*). Previous mammalian synthetic biology work has used as many as 12 unique genes in a single system (*34*). To scale up protein computation further will require expression of more starting protein species. It is likely that the major limitations are technical, relating to mammalian genome engineering, rather than problems with protein-protein interactions. We anticipate that as genome engineering techniques continue to advance (*35*), they will permit creation of larger systems, opening up the possibility of even more complex computation.

Finally, the perceptein output proteins are proteases, whose activities can be engineered to control diverse targets, including activation of endogenous pathways, such as cell death; transcription factors; or other synthetic protein systems (*21, 23*). They can also be wired to additional perceptein layers, allowing the construction of more powerful, and more fault-tolerant, multi-layer networks (*36*). Future iterations of this design could therefore enable the programming of more complex biochemical computations, rivaling those produced by evolution, in living cells.

## Acknowledgements

We thank L. Chong, X.J. Gao, and U. Alon for scientific input; J. Gregrowicz, R. Du, B. Emert, B. Gu, F. Horns, D. Li, A. Lu, K. Luo, Y. Takei, and S. Xia for critical feedback.

## Funding

This research was supported by the National Institute of Health grant R01 MH116508 and the Allen Discovery Center program under Award No. UWSC10142, a Paul G. Allen Frontiers Group advised program of the Paul G. Allen Family Foundation. M.B.E. is a Howard Hughes Medical Institute Investigator. Z.C. is supported by the Damon Runyon Cancer Research Foundation DRG-2388-20 and is a fellow of the Burroughs Wellcome Fund Career Awards at the Scientific Interface.

## Author contributions

Z.C. and M.B.E. conceived and designed the study; Z.C. performed mathematical modeling with help from R.Z. and M.B.E.; Z.C. and J.M.L. performed experiments; R.Z. constructed the reporter cell line; Z.C. and M.B.E. wrote the manuscript with input from all authors.

## Competing interests

Z.C. and M.B.E. have filed a provisional patent application based on this work.

## Data and materials availability

Raw data and code used for simulation and data analysis can be downloaded from https://data.caltech.edu/records/20215.

This article is subject to HHMI’s Open Access to Publications policy. HHMI lab heads have previously granted a nonexclusive CC BY 4.0 license to the public and a sublicensable license to HHMI in their research articles. Pursuant to those licenses, the author-accepted manuscript of this article can be made freely available under a CC BY 4.0 license immediately upon publication.

## Supplementary Materials

Materials and Methods

Supplementary Text

Figs. S1 to S3

Tables S1 to S3

## References and Notes

1. A. Hafner, M. L. Bulyk, A. Jambhekar, G. Lahav, The multiple mechanisms that regulate p53 activity and cell fate. Nat. Rev. Mol. Cell Biol. 20, 199–210 (2019).

2. M. Zagorski, Y. Tabata, N. Brandenberg, M. P. Lutolf, G. Tkacik, T. Bollenbach, J. Briscoe, Kicheva, Decoding of position in the developing neural tube from antiparallel morphogen gradients. Science. 356, 1379–1383 (2017).

3. Y. E. Antebi, S. Reich-Zeliger, Y. Hart, A. Mayo, I. Eizenberg, J. Rimer, P. Putheti, D. Pe’er, N. Friedman, Mapping differentiation under mixed culture conditions reveals a tunable continuum of T cell fates. PLoS Biol. 11, e1001616 (2013).

4. M. Hong, J. D. Clubb, Y. Y. Chen, Engineering CAR-T Cells for Next-Generation Cancer Therapy. Cancer Cell (2020), doi:10.1016/j.ccell.2020.07.005.

5. Y. Benenson, Biomolecular computing systems: principles, progress and potential. Nat. Rev. Genet. 13, 455–468 (2012).

6. W. Maass, On the computational power of winner-take-all. Neural Comput. 12, 2519–2535 (2000).

7. A. J. Genot, T. Fujii, Y. Rondelez, Computing with competition in biochemical networks. Phys. Rev. Lett. 109, 208102 (2012).

8. Genot Anthony J. Fujii Teruo, Rondelez Yannick, Scaling down DNA circuits with competitive neural networks. J. R. Soc. Interface. 10, 20130212 (2013).

9. C. C. Samaniego, A. Moorman, G. Giordano, E. Franco, Signaling-based neural networks for cellular computation. Cold Spring Harbor Laboratory (2020), p. 2020.11.10.377077.

10. O. Kanakov, R. Kotelnikov, A. Alsaedi, L. Tsimring, R. Huerta, A. Zaikin, M. Ivanchenko, Multi-input distributed classifiers for synthetic genetic circuits. PLoS One. 10, e0125144 (2015).

11. A. Didovyk, O. I. Kanakov, M. V. Ivanchenko, J. Hasty, R. Huerta, L. Tsimring, Distributed classifier based on genetically engineered bacterial cell cultures. ACS Synth. Biol. 4, 72–82 (2015).

12. L. Qian, E. Winfree, J. Bruck, Neural network computation with DNA strand displacement cascades. Nature. 475, 368–372 (2011).

13. K. M. Cherry, L. Qian, Scaling up molecular pattern recognition with DNA-based winner-take-all neural networks. Nature. 559, 370–376 (2018).

14. Z. Xie, L. Wroblewska, L. Prochazka, R. Weiss, Y. Benenson, Multi-input RNAi-based logic circuit for identification of specific cancer cells. Science. 333, 1307–1311 (2011).

15. P. Mohammadi, N. Beerenwinkel, Y. Benenson, Automated Design of Synthetic Cell Classifier Circuits Using a Two-Step Optimization Strategy. Cell Syst. 4, 207–218.e14 (2017).

16. Z. Chen, M. B. Elowitz, Programmable protein circuit design. Cell. 184, 2284–2301 (2021).

17. Z. Chen, S. E. Boyken, M. Jia, F. Busch, D. Flores-Solis, M. J. Bick, P. Lu, Z. L. VanAernum Sahasrabuddhe, R. A. Langan, S. Bermeo, T. J. Brunette, V. K. Mulligan, L. P. Carter, F. DiMaio, N. G. Sgourakis, V. H. Wysocki, D. Baker, Programmable design of orthogonal protein heterodimers. Nature. 565, 106–111 (2019).

18. F. Thomas, A. L. Boyle, A. J. Burton, D. N. Woolfson, A set of de novo designed parallel heterodimeric coiled coils with quantified dissociation constants in the micromolar to sub-nanomolar regime. J. Am. Chem. Soc. 135, 5161–5166 (2013).

19. H. Gradisar, R. Jerala, De novo design of orthogonal peptide pairs forming parallel coiled-coil heterodimers. J. Pept. Sci. 17, 100–106 (2011).

20. A. W. Reinke, R. A. Grant, A. E. Keating, A synthetic coiled-coil interactome provides heterospecific modules for molecular engineering. J. Am. Chem. Soc. 132, 6025–6031 (2010).

21. X. J. Gao, L. S. Chong, M. S. Kim, M. B. Elowitz, Programmable protein circuits in living cells. Science. 361, 1252–1258 (2018).

22. T. Fink, J. Lonzaric, A. Praznik, T. Plaper, E. Merljak, K. Leben, N. Jerala, T. Lebar, Z. Strmsek, F. Lapenta, M. Bencina, R. Jerala, Design of fast proteolysis-based signaling and logic circuits in mammalian cells. Nat. Chem. Biol. 15, 115–122 (2019).

23. H. K. Chung, X. Zou, B. T. Bajar, V. R. Brand, Y. Huo, J. F. Alcudia, J. E. Ferrell Jr, M. Z. Lin, A compact synthetic pathway rewires cancer signaling to therapeutic effector release. Science. 364 (2019), doi:10.1126/science.aat6982.

24. R. M. Hartfield, K. A. Schwarz, J. J. Muldoon, N. Bagheri, J. N. Leonard, Multiplexing Engineered Receptors for Multiparametric Evaluation of Environmental Ligands. ACS Synth. Biol. 6, 2042–2055 (2017).

25. J. Kim, J. Hopfield, E. Winfree, in Advances in Neural Information Processing Systems 17, L. K. Saul, Y. Weiss, L. Bottou, Eds. (MIT Press, 2005), pp. 681–688.

26. Z. Chen, R. D. Kibler, A. Hunt, F. Busch, J. Pearl, M. Jia, Z. L. VanAernum, B. I. M. Wicky, G. Dods, H. Liao, M. S. Wilken, C. Ciarlo, S. Green, H. El-Samad, J. Stamatoyannopoulos, V. H. Wysocki, M. C. Jewett, S. E. Boyken, D. Baker, De novo design of protein logic gates. Science. 368, 78–84 (2020).

27. J. Hertz, A. Krogh, R. G. Palmer, Introduction to the theory of neural computation (CRC Press, 2018).

28. R. Milo, P. Jorgensen, U. Moran, G. Weber, M. Springer, BioNumbers--the database of key numbers in molecular and cell biology. Nucleic Acids Res. 38, D750–3 (2010).

29. M. B. Elowitz, A. J. Levine, E. D. Siggia, P. S. Swain, Stochastic gene expression in a single cell. Science. 297, 1183–1186 (2002).

30. K. E. Prehoda, J. A. Scott, R. D. Mullins, W. A. Lim, Integration of multiple signals through cooperative regulation of the N-WASP-Arp2/3 complex. Science. 290, 801–806 (2000).

31. D. Schmucker, J. C. Clemens, H. Shu, C. A. Worby, J. Xiao, M. Muda, J. E. Dixon, S. L. Zipursky, Drosophila Dscam is an axon guidance receptor exhibiting extraordinary molecular diversity. Cell. 101, 671–684 (2000).

32. I. Nylander, K. Tan-No, A. Winter, J. Silberring, Processing of prodynorphin-derived peptides in striatal extracts. Identification by electrospray ionization mass spectrometry linked to size-exclusion chromatography. Life Sci. 57, 123–129 (1995).

33. B. L. Allen, D. J. Taatjes, The Mediator complex: a central integrator of transcription. Nat. Rev. Mol. Cell Biol. 16, 155–166 (2015).

34. A. Farhadi, G. H. Ho, D. P. Sawyer, R. W. Bourdeau, M. G. Shapiro, Ultrasound imaging of gene expression in mammalian cells. Science. 365, 1469–1475 (2019).

35. M. Zhang, C. Yang, I. Tasan, H. Zhao, Expanding the Potential of Mammalian Genome Engineering via Targeted DNA Integration. ACS Synth. Biol. 10, 429–446 (2021).

36. U. Alon, An introduction to systems biology: design principles of biological circuits (Chapman and Hall/CRC, 2006).

37. D. T. Gillespie, A general method for numerically simulating the stochastic time evolution of coupled chemical reactions. J. Comput. Phys. 22, 403–434 (1976).

38. J. Bezanson, A. Edelman, S. Karpinski, V. B. Shah, Julia: A Fresh Approach to Numerical Computing. SIAM Rev. 59, 65–98 (2017).

39. M. K. Doherty, D. E. Hammond, M. J. Clague, S. J. Gaskell, R. J. Beynon, Turnover of the human proteome: determination of protein intracellular stability by dynamic SILAC. J. Proteome Res. 8, 104–112 (2009).

40. A. Peth, J. A. Nathan, A. L. Goldberg, The ATP costs and time required to degrade ubiquitinated proteins by the 26 S proteasome. J. Biol. Chem. 288, 29215–29222 (2013).

41. J. Tozser, J. E. Tropea, S. Cherry, P. Bagossi, T. D. Copeland, A. Wlodawer, D. S. Waugh, Comparison of the substrate specificity of two potyvirus proteases. FEBS J. 272, 514–523 (2005).

42. K. Luby-Phelps, Cytoarchitecture and physical properties of cytoplasm: volume, viscosity, diffusion, intracellular surface area. Int. Rev. Cytol. 192, 189–221 (2000).

