## Supplementary material for "A synthetic protein-level neural network in mammalian cells": SI

This PDF file includes:

Materials and Methods
Supplementary Text
Figs. S1 to S3
Tables S1 to S3

### Materials and Methods

Construction of synthetic genes

Some constructs were generated using standard cloning procedures. The inserts were generated using PCR or gBlock synthesis (IDT), and were annealed by Gibson assembly with backbones that are linearized using restriction digestion. The rest of the constructs were designed by authors and synthesized by Genscript. mRNAs were ordered from TriLink BioTechnologies. A list of all constructs used in this study is included in Table 1.

Tissue culture

The monoclonal reporter cell line HEK1012 was generated using the PiggyBac Transposon system (Systems Biosciences) in Flp-In™ T-REx™ Human Embryonic Kidney 293 cell line (HEK293-TRex, Thermo). The plasmid with 3-phosphoglycerate kinase (PGK) promoter driving the expression of mCitrine and mCherry with N-end protease activatable degrons in PiggyBac backbone was co-transfected with a Super PiggyBac Transposase plasmid into HEK293-TRex cells. 24 hours after transfection, cells were transferred into a 6-well plate and selected with 400 µg/ml Zeocin for 9 days (split into Zeocin media every 3 days). The resulting polyclonal cells were then diluted at 1 cell/well into 96-well plates. After a week, wells with a single clone and positive mCitrine and mCherry fluorescence were labeled. The HEK1012 line was one of the clones that has medium mCitrine and mCherry expression from flow cytometry measurement. Cells were maintained in Eppendorf CellXpert cell culture incubators at 37°C with 5% $CO_2$. Cells were grown in media containing Dulbecco's Modified Eagle Medium (Gibco) supplemented with 10% Fetal Bovine Serum (Avantor), 1 mM sodium pyruvate (Gibco), 10 unit/ml penicillin (Gibco), 10 µg/ml streptomycin (Gibco), 2 mM L-glutamine (Gibco) and 0.1 mM MEM non-essential amino acids (Gibco).

Transient transfection of DNA into reporter cells

HEK1012 reporter cells were seeded in 24-well plates at a density of 0.05 - 0.1 x $10^6$ cells per well and cultured for 24 hours. Transient transfection was performed the following day using Lipofectamine 2000 (Thermo Fisher) following the manufacturer's protocol. 100 ng/mL doxycycline was added to the growth media whenever expression is needed from a CMV-TO promoter. Cells were changed into fresh growth media, with 100 ng/mL doxycycline, 24 hours after transfection, and were analyzed by flow cytometry after 24 hours.

Transient transfection of mRNA into reporter cells

HEK1012 reporter cells were seeded in 24-well plates at a density of 0.05 x $10^6$ cells per well and cultured for 24 hours. Transient transfection was performed the following day using the TransIT®-mRNA Transfection Kit (Mirus Bio) following the manufacturer's protocol. Cells were analyzed by flow cytometry after 24 hours.

Flow cytometry

Cells in 24 well plates were trypsinized with 40 µL of 0.05% trypsin-EDTA (Gibco) for 1 minute at room temperature, and subsequently resuspended in 100 µL of Hanks' Balanced Salt Solution (HBSS) containing 2.5mg/ml Bovine Serum Albumin (BSA), 1mM ethylenediaminetetraacetic acid (EDTA), and 4 units/ml DNase I (NEB). Cells were then filtered through a 40 μm cell strainer (Falcon™) or a 96-well plate cell strainer (Millipore) and analyzed by flow cytometry (CytoFLEX, Beckman Coulter).

Fluorescent signal quantification

Flow cytometry data was processed using the Cytoflow python package (https://github.com/cytoflow/cytoflow). Events collected from flow cytometry experiments were first gated based on forward vs. side scatter to select for cells, followed by gating based on scatter parameters, forward area vs forward height, to select for single cells. Data were then gated on fluorescence of the blue fluorescent protein (BFP), emission 450/45, co-transfection marker between 98 and 99.5 percentiles. Median values were taken from mCitrine, 525/40, or mCherry, 610/20, output signals.

Calculation of protease activities

Because the HEK1012 cell line is constitutively expressing protease repressible fluorescent proteins, normalized protease activities were calculated using (N-O)/(N-P), where N is observed fluorescence when cells were transfected with the negative control plasmids (protease halves), P is observed fluorescence when cells were transfected with the positive control plasmids (protease halves fused to DHD domains), and O is observed fluorescence in each experiment.

Deterministic simulation of the winner-take-all neural network

We performed numerical simulations in Python to gain insights for the behavior of the circuit. Four types of interactions were modeled: protein synthesis, protein binding, protease cleavage, and first-order protein degradation. Here we present the list of possible chemical reactions in the order in which they occur (Figure 2A).

- Protein synthesis: $X_i$, $N_{ij}D$, $C_k$ denote the DHD inputs, N-half proteases, and C-half proteases, respectively. In the 2-input, 2-output system, they comprise 8 species. The superscript D denotes an attached degron (from DHFR). The subscripts i and j denote the identity of the DHD and protease halves, respectively.

$$\varnothing \rightarrow X_i$$
$$\varnothing \rightarrow N_{ij}^D$$
$$\varnothing \rightarrow C_k$$

- Protein binding I: Here the DHD, N-half, and C-half protease bind cooperatively to reconstitute the protease complex $C_kX_iN_{ij}^D$ (Figure 2B). These complexes are active only when j = k, otherwise they represent inactive hybrids, containing mismatched halves of different proteases.

$$X_i + N_{ij}^D \rightarrow X_iN_{ij}^D$$
$$C_k + X_iN_{ij}^D \rightarrow C_kX_iN_{ij}^D$$

- Protease cleavage (i), self-activation: active proteases remove the DHFR tags off proteases of the same kind. $C_kX_iN_{ij}$ is a reconstituted protease without the DHFR degradation domain. It is an active protease when j = k. Otherwise, it represents an inactive hybrid containing protease halves from different proteases. $N_{ij}$ is the N-half protease without the DHFR degradation domain.

*Proteases cleave trimers:*

$$C_kX_iN_{ij}^D + C_nX_lN_{lm}^D \rightarrow C_kX_iN_{ij}^D + C_nX_lN_{lm} \ (j = k = m)$$
$$C_kX_iN_{ij} + C_nX_lN_{lm}^D \rightarrow C_kX_iN_{ij} + C_nX_lN_{lm} \ (j = k = m)$$

*Proteases cleave dimers:*

$$C_kX_iN_{ij}^D + X_lN_{lm}^D \rightarrow C_kX_iN_{ij}^D + X_lN_{lm} \ (j = k = m)$$
$$C_kX_iN_{ij} + X_lN_{lm}^D \rightarrow C_kX_iN_{ij} + X_lN_{lm} \ (j = k = m)$$

*Proteases cleave monomers:*

$$C_kX_iN_{ij}^D + N_{lm}^D \rightarrow C_kX_iN_{ij}^D + N_{lm} \ (j = k = m)$$
$$C_kX_iN_{ij} + N_{lm}^D \rightarrow C_kX_iN_{ij} + N_{lm} \ (j = k = m)$$

- Protease cleavage (ii), mutual inhibition: active proteases cleave the C-half protease of different kinds off their DHD domains. Here $C_n^E$ is a DHD domain without an attached C-half protease (E for empty). $C_n^EX_lN_{lm}^D$ is a protease complex with the DHFR tag and without the C-half protease. $C_n^EX_lN_{lm}$ is a protease complex without the DHFR tag and without the C-half protease.

*Proteases cleave monomers:*

$$C_kX_iN_{ij}^D + C_n \rightarrow C_kX_iN_{ij}^D + C_n^E \ (j = k \neq n)$$
$$C_kX_iN_{ij} + C_n \rightarrow C_kX_iN_{ij}^D + C_n^E \ (j = k \neq n)$$

*Proteases cleave trimers:*

$$C_kX_iN_{ij}^D + C_nX_lN_{lm}^D \rightarrow C_kX_iN_{ij}^D + C_n^EX_lN_{lm}^D \ (j = k \neq n)$$
$$C_kX_iN_{ij} + C_nX_lN_{lm}^D \rightarrow C_kX_iN_{ij} + C_n^EX_lN_{lm}^D \ (j = k \neq n)$$

*Proteases cleave trimers:*

$$C_kX_iN_{ij}^D + C_nX_lN_{lm} \rightarrow C_kX_iN_{ij}^D + C_n^EX_lN_{lm} \ (j = k \neq n)$$
$$C_kX_iN_{ij} + C_nX_lN_{lm} \rightarrow C_kX_iN_{ij} + C_n^EX_lN_{lm} \ (j = k \neq n)$$

- Protein binding (ii): newly formed proteins such as $N_{ij}$ and $C_k^E$ can now participate in binding. $C_k^EX_iN_{ij}^D$ is a protease complex with the DHFR tag and without the C-half protease. $C_k^EX_iN_{ij}$ is a protease complex without the DHFR tag and without the C-half protease. Both complexes are inactive due to the missing C-half protease. In contrast, $C_kX_iN_{ij}$ is an active protease when j = k.

$$X_i + N_{ij} \rightarrow X_iN_{ij}$$
$$X_iN_{ij} + C_k \rightarrow C_kX_iN_{ij}$$
$$X_iN_{ij} + C_k^E \rightarrow C_k^EX_iN_{ij}$$

$$X_iN_{ij}^D + C_k^E \rightarrow C_k^EX_iN_{ij}^D$$

- Protease cleavage (iii): $C_n^EX_lN_{lm}^D$ is a new substrate for protease cleavage.

$$C_kX_iN_{ij}^D + C_n^EX_lN_{lm}^D \rightarrow C_kX_iN_{ij}^D + C_n^EX_lN_{lm} \ (j = k = m)$$
$$C_kX_iN_{ij} + C_n^EX_lN_{lm}^D \rightarrow C_kX_iN_{ij} + C_n^EX_lN_{lm} \ (j = k = m)$$

- Protein degradation: every species with a superscript D, indicating it contains a degron, is assumed to undergo faster degradation than the corresponding species without the superscript D (lacking the degron).

Given the large number of chemical reactions in the system, we wrote a Python script that automatically and programmably generates these reactions and their corresponding ordinary differential equations (ODEs), using reaction rates provided in Table 1. This system of ODEs was then solved using the odeint solver in Python.

Stochastic simulation of the winner-take-all neural network

To perform stochastic simulations using the Gillespie algorithm (*37*), we first converted mass action rate constants (k) to stochastic rate constants (c) using the following formulae:

0th order reactions: $c = N_A*V*k$
1st order reactions: $c = k$
2nd order reactions: $c = k/(N_A*V)$

where $N_A$ is the Avogadro constant and V is the cell volume. The resulting list of chemical reactions were simulated in Julia (*38*).

**Supplementary Text**

Here we present the full chemical reaction network for a 2-input, 2-node comparator.

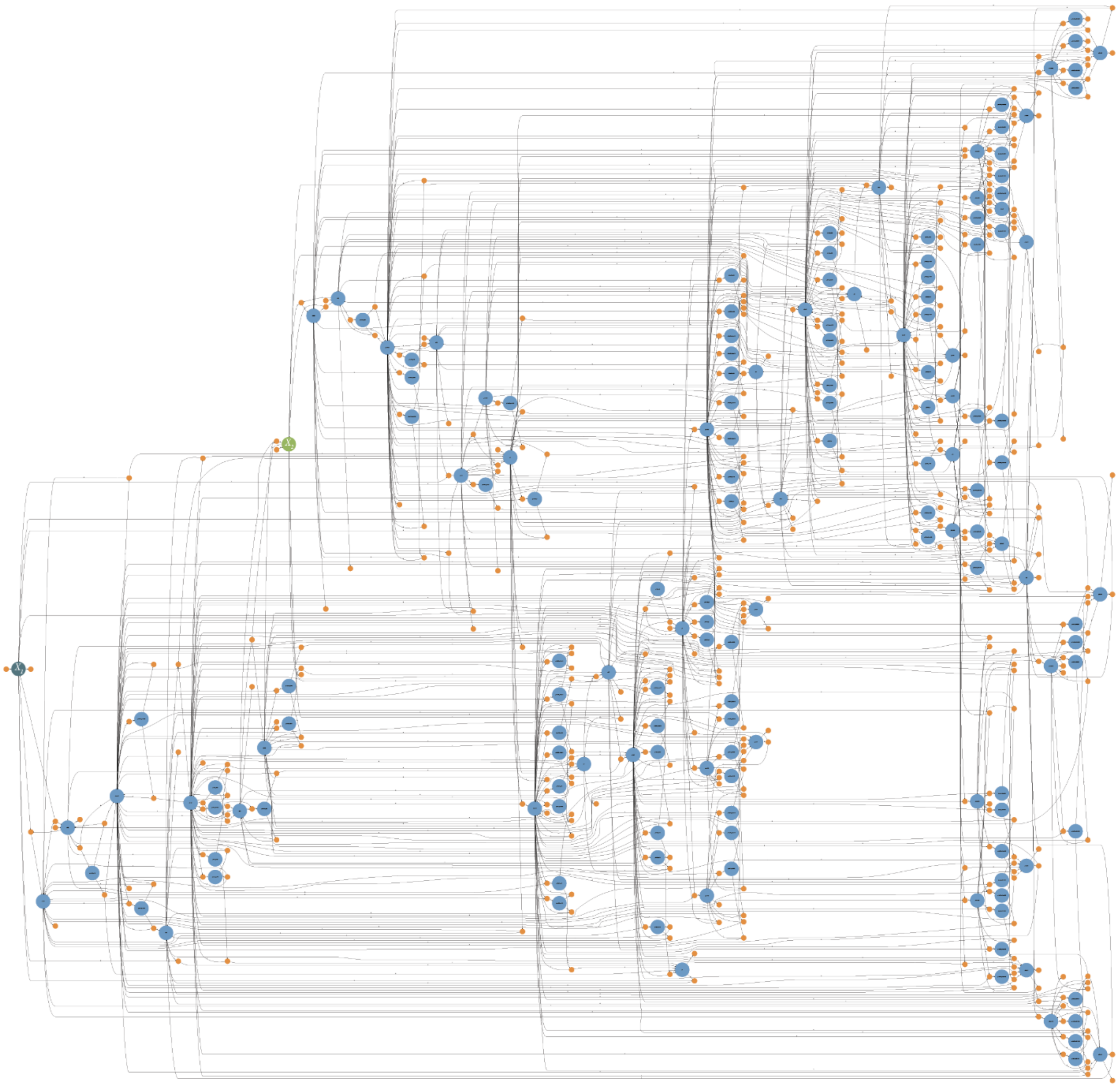

**A chemical reaction network of the 2-input comparator.** Small orange circles represent chemical reactions, and blue circles represent protein species. Black arrows from species to reactions indicate reactants, and are labeled with their input stoichiometry. Black arrows from reactions to species indicate formation of products, and are likewise labeled with their output stoichiometry. This graph was automatically generated using Julia.

In the page below we list all 310 chemical reactions that fully describe the system. “-” indicates an enzyme-substrate complex.

$$\mathrm{X_1 + N_{11}{}^D \underset{koff_1}{\overset{kon_1}{\rightleftharpoons}} X_1N_{11}{}^D} \tag{1}$$

$$\mathrm{X_1 + N_{12}{}^D \underset{koff_1}{\overset{kon_1}{\rightleftharpoons}} X_1N_{12}{}^D} \tag{2}$$

$$\mathrm{X_2 + N_{21}{}^D \underset{koff_1}{\overset{kon_1}{\rightleftharpoons}} X_2N_{21}{}^D} \tag{3}$$

$$\mathrm{X_2 + N_{22}{}^D \underset{koff_1}{\overset{kon_1}{\rightleftharpoons}} X_2N_{22}{}^D} \tag{4}$$

$$\mathrm{C_1 + X_1N_{11}{}^D \underset{koff_2}{\overset{kon_2}{\rightleftharpoons}} C_1X_1N_{11}{}^D} \tag{5}$$

$$\mathrm{C_1 + X_1N_{12}{}^D \underset{koff_2}{\overset{kon_2}{\rightleftharpoons}} C_1X_1N_{12}{}^D} \tag{6}$$

$$\mathrm{C_1 + X_2N_{21}{}^D \underset{koff_2}{\overset{kon_2}{\rightleftharpoons}} C_1X_2N_{21}{}^D} \tag{7}$$

$$\mathrm{C_1 + X_2N_{22}{}^D \underset{koff_2}{\overset{kon_2}{\rightleftharpoons}} C_1X_2N_{22}{}^D} \tag{8}$$

$$\mathrm{C_2 + X_1N_{11}{}^D \underset{koff_2}{\overset{kon_2}{\rightleftharpoons}} C_2X_1N_{11}{}^D} \tag{9}$$

$$\mathrm{C_2 + X_1N_{12}{}^D \underset{koff_2}{\overset{kon_2}{\rightleftharpoons}} C_2X_1N_{12}{}^D} \tag{10}$$

$$\mathrm{C_2 + X_2N_{21}{}^D \underset{koff_2}{\overset{kon_2}{\rightleftharpoons}} C_2X_2N_{21}{}^D} \tag{11}$$

$$\mathrm{C_2 + X_2N_{22}{}^D \underset{koff_2}{\overset{kon_2}{\rightleftharpoons}} C_2X_2N_{22}{}^D} \tag{12}$$

$$\mathrm{C_1X_1N_{11}{}^D + N_{11}{}^D \underset{koff_{\mathit{p}1}}{\overset{kon_{\mathit{p}1}}{\rightleftharpoons}} C_1X_1N_{11}{}^D{-}N_{11}{}^D} \tag{13}$$

$$\mathrm{C_1X_1N_{11}{}^D + N_{21}{}^D \underset{koff_{\mathit{p}1}}{\overset{kon_{\mathit{p}1}}{\rightleftharpoons}} C_1X_1N_{11}{}^D{-}N_{21}{}^D} \tag{14}$$

$$\mathrm{C_1X_2N_{21}{}^D + N_{11}{}^D \underset{koff_{\mathit{p}1}}{\overset{kon_{\mathit{p}1}}{\rightleftharpoons}} C_1X_2N_{21}{}^D{-}N_{11}{}^D} \tag{15}$$

$$\mathrm{C_1X_2N_{21}{}^D + N_{21}{}^D \underset{koff_{\mathit{p}1}}{\overset{kon_{\mathit{p}1}}{\rightleftharpoons}} C_1X_2N_{21}{}^D{-}N_{21}{}^D} \tag{16}$$

$$\mathrm{C_2X_1N_{12}{}^D + N_{12}{}^D \underset{koff_{\mathit{p}2}}{\overset{kon_{\mathit{p}2}}{\rightleftharpoons}} C_2X_1N_{12}{}^D{-}N_{12}{}^D} \tag{17}$$

$$\mathrm{C_2X_1N_{12}{}^D + N_{22}{}^D \underset{koff_{\mathit{p}2}}{\overset{kon_{\mathit{p}2}}{\rightleftharpoons}} C_2X_1N_{12}{}^D{-}N_{22}{}^D} \tag{18}$$

$$\mathrm{C_2X_2N_{22}{}^D + N_{12}{}^D \underset{koff_{\mathit{p}2}}{\overset{kon_{\mathit{p}2}}{\rightleftharpoons}} C_2X_2N_{22}{}^D{-}N_{12}{}^D} \tag{19}$$

$$\mathrm{C_2X_2N_{22}{}^D + N_{22}{}^D \underset{koff_{\mathit{p}2}}{\overset{kon_{\mathit{p}2}}{\rightleftharpoons}} C_2X_2N_{22}{}^D{-}N_{22}{}^D} \tag{20}$$

$$\mathrm{C_1X_1N_{11}{}^D{-}N_{11}{}^D \xrightarrow{kcat_1} C_1X_1N_{11}{}^D + N_{11}} \tag{21}$$

$$\mathrm{C_1X_1N_{11}{}^D{-}N_{21}{}^D \xrightarrow{kcat_1} C_1X_1N_{11}{}^D + N_{21}} \tag{22}$$

$$\mathrm{C_1X_2N_{21}{}^D{-}N_{11}{}^D \xrightarrow{kcat_1} C_1X_2N_{21}{}^D + N_{11}} \tag{23}$$

$$\mathrm{C_1X_2N_{21}{}^D{-}N_{21}{}^D \xrightarrow{kcat_1} C_1X_2N_{21}{}^D + N_{21}} \tag{24}$$

$$\mathrm{C_2X_1N_{12}{}^D{-}N_{12}{}^D \xrightarrow{kcat_2} C_2X_1N_{12}{}^D + N_{12}} \tag{25}$$

$$C_2X_1N_{12}{}^{D}{-}N_{22}{}^{D} \xrightarrow{kcat_2} C_2X_1N_{12}{}^{D} + N_{22} \tag{26}$$

$$C_2X_2N_{22}{}^{D}{-}N_{12}{}^{D} \xrightarrow{kcat_2} C_2X_2N_{22}{}^{D} + N_{12} \tag{27}$$

$$C_2X_2N_{22}{}^{D}{-}N_{22}{}^{D} \xrightarrow{kcat_2} C_2X_2N_{22}{}^{D} + N_{22} \tag{28}$$

$$C_1X_1N_{11}{}^{D} + X_1N_{11}{}^{D} \underset{koff_{p1}}{\overset{kon_{p1}}{\rightleftharpoons}} C_1X_1N_{11}{}^{D}{-}X_1N_{11}{}^{D} \tag{29}$$

$$C_1X_1N_{11}{}^{D} + X_2N_{21}{}^{D} \underset{koff_{p1}}{\overset{kon_{p1}}{\rightleftharpoons}} C_1X_1N_{11}{}^{D}{-}X_2N_{21}{}^{D} \tag{30}$$

$$C_1X_2N_{21}{}^{D} + X_1N_{11}{}^{D} \underset{koff_{p1}}{\overset{kon_{p1}}{\rightleftharpoons}} C_1X_2N_{21}{}^{D}{-}X_1N_{11}{}^{D} \tag{31}$$

$$C_1X_2N_{21}{}^{D} + X_2N_{21}{}^{D} \underset{koff_{p1}}{\overset{kon_{p1}}{\rightleftharpoons}} C_1X_2N_{21}{}^{D}{-}X_2N_{21}{}^{D} \tag{32}$$

$$C_2X_1N_{12}{}^{D} + X_1N_{12}{}^{D} \underset{koff_{p2}}{\overset{kon_{p2}}{\rightleftharpoons}} C_2X_1N_{12}{}^{D}{-}X_1N_{12}{}^{D} \tag{33}$$

$$C_2X_1N_{12}{}^{D} + X_2N_{22}{}^{D} \underset{koff_{p2}}{\overset{kon_{p2}}{\rightleftharpoons}} C_2X_1N_{12}{}^{D}{-}X_2N_{22}{}^{D} \tag{34}$$

$$C_2X_2N_{22}{}^{D} + X_1N_{12}{}^{D} \underset{koff_{p2}}{\overset{kon_{p2}}{\rightleftharpoons}} C_2X_2N_{22}{}^{D}{-}X_1N_{12}{}^{D} \tag{35}$$

$$C_2X_2N_{22}{}^{D} + X_2N_{22}{}^{D} \underset{koff_{p2}}{\overset{kon_{p2}}{\rightleftharpoons}} C_2X_2N_{22}{}^{D}{-}X_2N_{22}{}^{D} \tag{36}$$

$$C_1X_1N_{11}{}^{D}{-}X_1N_{11}{}^{D} \xrightarrow{kcat_1} C_1X_1N_{11}{}^{D} + X_1N_{11} \tag{37}$$

$$C_1X_1N_{11}{}^{D}{-}X_2N_{21}{}^{D} \xrightarrow{kcat_1} C_1X_1N_{11}{}^{D} + X_2N_{21} \tag{38}$$

$$C_1X_2N_{21}{}^{D}{-}X_1N_{11}{}^{D} \xrightarrow{kcat_1} C_1X_2N_{21}{}^{D} + X_1N_{11} \tag{39}$$

$$C_1X_2N_{21}{}^{D}{-}X_2N_{21}{}^{D} \xrightarrow{kcat_1} C_1X_2N_{21}{}^{D} + X_2N_{21} \tag{40}$$

$$C_2X_1N_{12}{}^{D}{-}X_1N_{12}{}^{D} \xrightarrow{kcat_2} C_2X_1N_{12}{}^{D} + X_1N_{12} \tag{41}$$

$$C_2X_1N_{12}{}^{D}{-}X_2N_{22}{}^{D} \xrightarrow{kcat_2} C_2X_1N_{12}{}^{D} + X_2N_{22} \tag{42}$$

$$C_2X_2N_{22}{}^{D}{-}X_1N_{12}{}^{D} \xrightarrow{kcat_2} C_2X_2N_{22}{}^{D} + X_1N_{12} \tag{43}$$

$$C_2X_2N_{22}{}^{D}{-}X_2N_{22}{}^{D} \xrightarrow{kcat_2} C_2X_2N_{22}{}^{D} + X_2N_{22} \tag{44}$$

$$2\,C_1X_1N_{11}{}^{D} \underset{koff_{p1}}{\overset{kon_{p1}}{\rightleftharpoons}} C_1X_1N_{11}{}^{D}{-}C_1X_1N_{11}{}^{D} \tag{45}$$

$$C_1X_1N_{11}{}^{D} + C_1X_2N_{21}{}^{D} \underset{koff_{p1}}{\overset{kon_{p1}}{\rightleftharpoons}} C_1X_1N_{11}{}^{D}{-}C_1X_2N_{21}{}^{D} \tag{46}$$

$$C_1X_1N_{11}{}^{D} + C_2X_1N_{11}{}^{D} \underset{koff_{p1}}{\overset{kon_{p1}}{\rightleftharpoons}} C_1X_1N_{11}{}^{D}{-}C_2X_1N_{11}{}^{D} \tag{47}$$

$$C_1X_1N_{11}{}^{D} + C_2X_2N_{21}{}^{D} \underset{koff_{p1}}{\overset{kon_{p1}}{\rightleftharpoons}} C_1X_1N_{11}{}^{D}{-}C_2X_2N_{21}{}^{D} \tag{48}$$

$$C_1X_2N_{21}{}^{D} + C_1X_1N_{11}{}^{D} \underset{koff_{p1}}{\overset{kon_{p1}}{\rightleftharpoons}} C_1X_2N_{21}{}^{D}{-}C_1X_1N_{11}{}^{D} \tag{49}$$

$$2\,C_1X_2N_{21}{}^{D} \underset{koff_{p1}}{\overset{kon_{p1}}{\rightleftharpoons}} C_1X_2N_{21}{}^{D}{-}C_1X_2N_{21}{}^{D} \tag{50}$$

$$\mathrm{C_1X_2N_{21}}^{\mathrm{D}} + \mathrm{C_2X_1N_{11}}^{\mathrm{D}} \xrightleftharpoons[\mathrm{koff}_{p1}]{\mathrm{kon}_{p1}} \mathrm{C_1X_2N_{21}}^{\mathrm{D}}{-}\mathrm{C_2X_1N_{11}}^{\mathrm{D}} \tag{51}$$

$$\mathrm{C_1X_2N_{21}}^{\mathrm{D}} + \mathrm{C_2X_2N_{21}}^{\mathrm{D}} \xrightleftharpoons[\mathrm{koff}_{p1}]{\mathrm{kon}_{p1}} \mathrm{C_1X_2N_{21}}^{\mathrm{D}}{-}\mathrm{C_2X_2N_{21}}^{\mathrm{D}} \tag{52}$$

$$\mathrm{C_2X_1N_{12}}^{\mathrm{D}} + \mathrm{C_1X_1N_{12}}^{\mathrm{D}} \xrightleftharpoons[\mathrm{koff}_{p2}]{\mathrm{kon}_{p2}} \mathrm{C_2X_1N_{12}}^{\mathrm{D}}{-}\mathrm{C_1X_1N_{12}}^{\mathrm{D}} \tag{53}$$

$$\mathrm{C_2X_1N_{12}}^{\mathrm{D}} + \mathrm{C_1X_2N_{22}}^{\mathrm{D}} \xrightleftharpoons[\mathrm{koff}_{p2}]{\mathrm{kon}_{p2}} \mathrm{C_2X_1N_{12}}^{\mathrm{D}}{-}\mathrm{C_1X_2N_{22}}^{\mathrm{D}} \tag{54}$$

$$2\,\mathrm{C_2X_1N_{12}}^{\mathrm{D}} \xrightleftharpoons[\mathrm{koff}_{p2}]{\mathrm{kon}_{p2}} \mathrm{C_2X_1N_{12}}^{\mathrm{D}}{-}\mathrm{C_2X_1N_{12}}^{\mathrm{D}} \tag{55}$$

$$\mathrm{C_2X_1N_{12}}^{\mathrm{D}} + \mathrm{C_2X_2N_{22}}^{\mathrm{D}} \xrightleftharpoons[\mathrm{koff}_{p2}]{\mathrm{kon}_{p2}} \mathrm{C_2X_1N_{12}}^{\mathrm{D}}{-}\mathrm{C_2X_2N_{22}}^{\mathrm{D}} \tag{56}$$

$$\mathrm{C_2X_2N_{22}}^{\mathrm{D}} + \mathrm{C_1X_1N_{12}}^{\mathrm{D}} \xrightleftharpoons[\mathrm{koff}_{p2}]{\mathrm{kon}_{p2}} \mathrm{C_2X_2N_{22}}^{\mathrm{D}}{-}\mathrm{C_1X_1N_{12}}^{\mathrm{D}} \tag{57}$$

$$\mathrm{C_2X_2N_{22}}^{\mathrm{D}} + \mathrm{C_1X_2N_{22}}^{\mathrm{D}} \xrightleftharpoons[\mathrm{koff}_{p2}]{\mathrm{kon}_{p2}} \mathrm{C_2X_2N_{22}}^{\mathrm{D}}{-}\mathrm{C_1X_2N_{22}}^{\mathrm{D}} \tag{58}$$

$$\mathrm{C_2X_2N_{22}}^{\mathrm{D}} + \mathrm{C_2X_1N_{12}}^{\mathrm{D}} \xrightleftharpoons[\mathrm{koff}_{p2}]{\mathrm{kon}_{p2}} \mathrm{C_2X_2N_{22}}^{\mathrm{D}}{-}\mathrm{C_2X_1N_{12}}^{\mathrm{D}} \tag{59}$$

$$2\,\mathrm{C_2X_2N_{22}}^{\mathrm{D}} \xrightleftharpoons[\mathrm{koff}_{p2}]{\mathrm{kon}_{p2}} \mathrm{C_2X_2N_{22}}^{\mathrm{D}}{-}\mathrm{C_2X_2N_{22}}^{\mathrm{D}} \tag{60}$$

$$\mathrm{C_1X_1N_{11}}^{\mathrm{D}}{-}\mathrm{C_1X_1N_{11}}^{\mathrm{D}} \xrightarrow{\mathrm{kcat}_1} \mathrm{C_1X_1N_{11}}^{\mathrm{D}} + \mathrm{C_1X_1N_{11}} \tag{61}$$

$$\mathrm{C_1X_1N_{11}}^{\mathrm{D}}{-}\mathrm{C_1X_2N_{21}}^{\mathrm{D}} \xrightarrow{\mathrm{kcat}_1} \mathrm{C_1X_1N_{11}}^{\mathrm{D}} + \mathrm{C_1X_2N_{21}} \tag{62}$$

$$\mathrm{C_1X_1N_{11}}^{\mathrm{D}}{-}\mathrm{C_2X_1N_{11}}^{\mathrm{D}} \xrightarrow{\mathrm{kcat}_1} \mathrm{C_1X_1N_{11}}^{\mathrm{D}} + \mathrm{C_2X_1N_{11}} \tag{63}$$

$$\mathrm{C_1X_1N_{11}}^{\mathrm{D}}{-}\mathrm{C_2X_2N_{21}}^{\mathrm{D}} \xrightarrow{\mathrm{kcat}_1} \mathrm{C_1X_1N_{11}}^{\mathrm{D}} + \mathrm{C_2X_2N_{21}} \tag{64}$$

$$\mathrm{C_1X_2N_{21}}^{\mathrm{D}}{-}\mathrm{C_1X_1N_{11}}^{\mathrm{D}} \xrightarrow{\mathrm{kcat}_1} \mathrm{C_1X_2N_{21}}^{\mathrm{D}} + \mathrm{C_1X_1N_{11}} \tag{65}$$

$$\mathrm{C_1X_2N_{21}}^{\mathrm{D}}{-}\mathrm{C_1X_2N_{21}}^{\mathrm{D}} \xrightarrow{\mathrm{kcat}_1} \mathrm{C_1X_2N_{21}}^{\mathrm{D}} + \mathrm{C_1X_2N_{21}} \tag{66}$$

$$\mathrm{C_1X_2N_{21}}^{\mathrm{D}}{-}\mathrm{C_2X_1N_{11}}^{\mathrm{D}} \xrightarrow{\mathrm{kcat}_1} \mathrm{C_1X_2N_{21}}^{\mathrm{D}} + \mathrm{C_2X_1N_{11}} \tag{67}$$

$$\mathrm{C_1X_2N_{21}}^{\mathrm{D}}{-}\mathrm{C_2X_2N_{21}}^{\mathrm{D}} \xrightarrow{\mathrm{kcat}_1} \mathrm{C_1X_2N_{21}}^{\mathrm{D}} + \mathrm{C_2X_2N_{21}} \tag{68}$$

$$\mathrm{C_2X_1N_{12}}^{\mathrm{D}}{-}\mathrm{C_1X_1N_{12}}^{\mathrm{D}} \xrightarrow{\mathrm{kcat}_2} \mathrm{C_2X_1N_{12}}^{\mathrm{D}} + \mathrm{C_1X_1N_{12}} \tag{69}$$

$$\mathrm{C_2X_1N_{12}}^{\mathrm{D}}{-}\mathrm{C_1X_2N_{22}}^{\mathrm{D}} \xrightarrow{\mathrm{kcat}_2} \mathrm{C_2X_1N_{12}}^{\mathrm{D}} + \mathrm{C_1X_2N_{22}} \tag{70}$$

$$\mathrm{C_2X_1N_{12}}^{\mathrm{D}}{-}\mathrm{C_2X_1N_{12}}^{\mathrm{D}} \xrightarrow{\mathrm{kcat}_2} \mathrm{C_2X_1N_{12}}^{\mathrm{D}} + \mathrm{C_2X_1N_{12}} \tag{71}$$

$$\mathrm{C_2X_1N_{12}}^{\mathrm{D}}{-}\mathrm{C_2X_2N_{22}}^{\mathrm{D}} \xrightarrow{\mathrm{kcat}_2} \mathrm{C_2X_1N_{12}}^{\mathrm{D}} + \mathrm{C_2X_2N_{22}} \tag{72}$$

$$\mathrm{C_2X_2N_{22}}^{\mathrm{D}}{-}\mathrm{C_1X_1N_{12}}^{\mathrm{D}} \xrightarrow{\mathrm{kcat}_2} \mathrm{C_2X_2N_{22}}^{\mathrm{D}} + \mathrm{C_1X_1N_{12}} \tag{73}$$

$$\mathrm{C_2X_2N_{22}}^{\mathrm{D}}{-}\mathrm{C_1X_2N_{22}}^{\mathrm{D}} \xrightarrow{\mathrm{kcat}_2} \mathrm{C_2X_2N_{22}}^{\mathrm{D}} + \mathrm{C_1X_2N_{22}} \tag{74}$$

$$\mathrm{C_2X_2N_{22}}^{\mathrm{D}}{-}\mathrm{C_2X_1N_{12}}^{\mathrm{D}} \xrightarrow{\mathrm{kcat}_2} \mathrm{C_2X_2N_{22}}^{\mathrm{D}} + \mathrm{C_2X_1N_{12}} \tag{75}$$

$$C_2X_2{N_{22}}^D{-}C_2X_2{N_{22}}^D \xrightarrow{kcat_2} C_2X_2{N_{22}}^D + C_2X_2N_{22} \tag{76}$$

$$C_1X_1{N_{11}}^D + C_2 \underset{koff_{p1}}{\overset{kon_{p1}}{\rightleftharpoons}} C_1X_1{N_{11}}^D{-}C_2 \tag{77}$$

$$C_1X_2{N_{21}}^D + C_2 \underset{koff_{p1}}{\overset{kon_{p1}}{\rightleftharpoons}} C_1X_2{N_{21}}^D{-}C_2 \tag{78}$$

$$C_2X_1{N_{12}}^D + C_1 \underset{koff_{p2}}{\overset{kon_{p2}}{\rightleftharpoons}} C_2X_1{N_{12}}^D{-}C_1 \tag{79}$$

$$C_2X_2{N_{22}}^D + C_1 \underset{koff_{p2}}{\overset{kon_{p2}}{\rightleftharpoons}} C_2X_2{N_{22}}^D{-}C_1 \tag{80}$$

$$C_1X_1{N_{11}}^D{-}C_2 \xrightarrow{kcat_1} C_1X_1{N_{11}}^D + {C_2}^E \tag{81}$$

$$C_1X_2{N_{21}}^D{-}C_2 \xrightarrow{kcat_1} C_1X_2{N_{21}}^D + {C_2}^E \tag{82}$$

$$C_2X_1{N_{12}}^D{-}C_1 \xrightarrow{kcat_2} C_2X_1{N_{12}}^D + {C_1}^E \tag{83}$$

$$C_2X_2{N_{22}}^D{-}C_1 \xrightarrow{kcat_2} C_2X_2{N_{22}}^D + {C_1}^E \tag{84}$$

$$X_1 + N_{11} \underset{koff_1}{\overset{kon_1}{\rightleftharpoons}} X_1N_{11} \tag{85}$$

$$X_1 + N_{12} \underset{koff_1}{\overset{kon_1}{\rightleftharpoons}} X_1N_{12} \tag{86}$$

$$X_2 + N_{21} \underset{koff_1}{\overset{kon_1}{\rightleftharpoons}} X_2N_{21} \tag{87}$$

$$X_2 + N_{22} \underset{koff_1}{\overset{kon_1}{\rightleftharpoons}} X_2N_{22} \tag{88}$$

$$C_1 + X_1N_{11} \underset{koff_2}{\overset{kon_2}{\rightleftharpoons}} C_1X_1N_{11} \tag{89}$$

$$C_1 + X_1N_{12} \underset{koff_2}{\overset{kon_2}{\rightleftharpoons}} C_1X_1N_{12} \tag{90}$$

$$C_1 + X_2N_{21} \underset{koff_2}{\overset{kon_2}{\rightleftharpoons}} C_1X_2N_{21} \tag{91}$$

$$C_1 + X_2N_{22} \underset{koff_2}{\overset{kon_2}{\rightleftharpoons}} C_1X_2N_{22} \tag{92}$$

$$C_2 + X_1N_{11} \underset{koff_2}{\overset{kon_2}{\rightleftharpoons}} C_2X_1N_{11} \tag{93}$$

$$C_2 + X_1N_{12} \underset{koff_2}{\overset{kon_2}{\rightleftharpoons}} C_2X_1N_{12} \tag{94}$$

$$C_2 + X_2N_{21} \underset{koff_2}{\overset{kon_2}{\rightleftharpoons}} C_2X_2N_{21} \tag{95}$$

$$C_2 + X_2N_{22} \underset{koff_2}{\overset{kon_2}{\rightleftharpoons}} C_2X_2N_{22} \tag{96}$$

$$C_1X_1N_{11} + {N_{11}}^D \underset{koff_{p1}}{\overset{kon_{p1}}{\rightleftharpoons}} C_1X_1N_{11}{-}{N_{11}}^D \tag{97}$$

$$C_1X_1N_{11} + {N_{21}}^D \underset{koff_{p1}}{\overset{kon_{p1}}{\rightleftharpoons}} C_1X_1N_{11}{-}{N_{21}}^D \tag{98}$$

$$C_1X_2N_{21} + {N_{11}}^D \underset{koff_{p1}}{\overset{kon_{p1}}{\rightleftharpoons}} C_1X_2N_{21}{-}{N_{11}}^D \tag{99}$$

$$C_1X_2N_{21} + {N_{21}}^D \underset{koff_{p1}}{\overset{kon_{p1}}{\rightleftharpoons}} C_1X_2N_{21}{-}{N_{21}}^D \tag{100}$$

$$C_2X_1N_{12} + N_{12}{}^{D} \underset{koff_{p2}}{\overset{kon_{p2}}{\rightleftharpoons}} C_2X_1N_{12}{-}N_{12}{}^{D} \tag{101}$$

$$C_2X_1N_{12} + N_{22}{}^{D} \underset{koff_{p2}}{\overset{kon_{p2}}{\rightleftharpoons}} C_2X_1N_{12}{-}N_{22}{}^{D} \tag{102}$$

$$C_2X_2N_{22} + N_{12}{}^{D} \underset{koff_{p2}}{\overset{kon_{p2}}{\rightleftharpoons}} C_2X_2N_{22}{-}N_{12}{}^{D} \tag{103}$$

$$C_2X_2N_{22} + N_{22}{}^{D} \underset{koff_{p2}}{\overset{kon_{p2}}{\rightleftharpoons}} C_2X_2N_{22}{-}N_{22}{}^{D} \tag{104}$$

$$C_1X_1N_{11}{-}N_{11}{}^{D} \xrightarrow{kcat_1} C_1X_1N_{11} + N_{11} \tag{105}$$

$$C_1X_1N_{11}{-}N_{21}{}^{D} \xrightarrow{kcat_1} C_1X_1N_{11} + N_{21} \tag{106}$$

$$C_1X_2N_{21}{-}N_{11}{}^{D} \xrightarrow{kcat_1} C_1X_2N_{21} + N_{11} \tag{107}$$

$$C_1X_2N_{21}{-}N_{21}{}^{D} \xrightarrow{kcat_1} C_1X_2N_{21} + N_{21} \tag{108}$$

$$C_2X_1N_{12}{-}N_{12}{}^{D} \xrightarrow{kcat_2} C_2X_1N_{12} + N_{12} \tag{109}$$

$$C_2X_1N_{12}{-}N_{22}{}^{D} \xrightarrow{kcat_2} C_2X_1N_{12} + N_{22} \tag{110}$$

$$C_2X_2N_{22}{-}N_{12}{}^{D} \xrightarrow{kcat_2} C_2X_2N_{22} + N_{12} \tag{111}$$

$$C_2X_2N_{22}{-}N_{22}{}^{D} \xrightarrow{kcat_2} C_2X_2N_{22} + N_{22} \tag{112}$$

$$C_1X_1N_{11} + X_1N_{11}{}^{D} \underset{koff_{p1}}{\overset{kon_{p1}}{\rightleftharpoons}} C_1X_1N_{11}{-}X_1N_{11}{}^{D} \tag{113}$$

$$C_1X_1N_{11} + X_2N_{21}{}^{D} \underset{koff_{p1}}{\overset{kon_{p1}}{\rightleftharpoons}} C_1X_1N_{11}{-}X_2N_{21}{}^{D} \tag{114}$$

$$C_1X_2N_{21} + X_1N_{11}{}^{D} \underset{koff_{p1}}{\overset{kon_{p1}}{\rightleftharpoons}} C_1X_2N_{21}{-}X_1N_{11}{}^{D} \tag{115}$$

$$C_1X_2N_{21} + X_2N_{21}{}^{D} \underset{koff_{p1}}{\overset{kon_{p1}}{\rightleftharpoons}} C_1X_2N_{21}{-}X_2N_{21}{}^{D} \tag{116}$$

$$C_2X_1N_{12} + X_1N_{12}{}^{D} \underset{koff_{p2}}{\overset{kon_{p2}}{\rightleftharpoons}} C_2X_1N_{12}{-}X_1N_{12}{}^{D} \tag{117}$$

$$C_2X_1N_{12} + X_2N_{22}{}^{D} \underset{koff_{p2}}{\overset{kon_{p2}}{\rightleftharpoons}} C_2X_1N_{12}{-}X_2N_{22}{}^{D} \tag{118}$$

$$C_2X_2N_{22} + X_1N_{12}{}^{D} \underset{koff_{p2}}{\overset{kon_{p2}}{\rightleftharpoons}} C_2X_2N_{22}{-}X_1N_{12}{}^{D} \tag{119}$$

$$C_2X_2N_{22} + X_2N_{22}{}^{D} \underset{koff_{p2}}{\overset{kon_{p2}}{\rightleftharpoons}} C_2X_2N_{22}{-}X_2N_{22}{}^{D} \tag{120}$$

$$C_1X_1N_{11}{-}X_1N_{11}{}^{D} \xrightarrow{kcat_1} C_1X_1N_{11} + X_1N_{11} \tag{121}$$

$$C_1X_1N_{11}{-}X_2N_{21}{}^{D} \xrightarrow{kcat_1} C_1X_1N_{11} + X_2N_{21} \tag{122}$$

$$C_1X_2N_{21}{-}X_1N_{11}{}^{D} \xrightarrow{kcat_1} C_1X_2N_{21} + X_1N_{11} \tag{123}$$

$$C_1X_2N_{21}{-}X_2N_{21}{}^{D} \xrightarrow{kcat_1} C_1X_2N_{21} + X_2N_{21} \tag{124}$$

$$C_2X_1N_{12}{-}X_1N_{12}{}^{D} \xrightarrow{kcat_2} C_2X_1N_{12} + X_1N_{12} \tag{125}$$

$$C_2X_1N_{12}{-}X_2N_{22}{}^{D} \xrightarrow{kcat_2} C_2X_1N_{12} + X_2N_{22} \tag{126}$$

$$C_2X_2N_{22}{-}X_1N_{12}{}^{D} \xrightarrow{kcat_2} C_2X_2N_{22} + X_1N_{12} \tag{127}$$

$$C_2X_2N_{22}{-}X_2N_{22}{}^{D} \xrightarrow{kcat_2} C_2X_2N_{22} + X_2N_{22} \tag{128}$$

$$C_1X_1N_{11} + C_1X_1N_{11}{}^{D} \underset{koff_{p1}}{\overset{kon_{p1}}{\rightleftharpoons}} C_1X_1N_{11}{-}C_1X_1N_{11}{}^{D} \tag{129}$$

$$C_1X_1N_{11} + C_1X_2N_{21}{}^{D} \underset{koff_{p1}}{\overset{kon_{p1}}{\rightleftharpoons}} C_1X_1N_{11}{-}C_1X_2N_{21}{}^{D} \tag{130}$$

$$C_1X_1N_{11} + C_2X_1N_{11}{}^{D} \underset{koff_{p1}}{\overset{kon_{p1}}{\rightleftharpoons}} C_1X_1N_{11}{-}C_2X_1N_{11}{}^{D} \tag{131}$$

$$C_1X_1N_{11} + C_2X_2N_{21}{}^{D} \underset{koff_{p1}}{\overset{kon_{p1}}{\rightleftharpoons}} C_1X_1N_{11}{-}C_2X_2N_{21}{}^{D} \tag{132}$$

$$C_1X_2N_{21} + C_1X_1N_{11}{}^{D} \underset{koff_{p1}}{\overset{kon_{p1}}{\rightleftharpoons}} C_1X_2N_{21}{-}C_1X_1N_{11}{}^{D} \tag{133}$$

$$C_1X_2N_{21} + C_1X_2N_{21}{}^{D} \underset{koff_{p1}}{\overset{kon_{p1}}{\rightleftharpoons}} C_1X_2N_{21}{-}C_1X_2N_{21}{}^{D} \tag{134}$$

$$C_1X_2N_{21} + C_2X_1N_{11}{}^{D} \underset{koff_{p1}}{\overset{kon_{p1}}{\rightleftharpoons}} C_1X_2N_{21}{-}C_2X_1N_{11}{}^{D} \tag{135}$$

$$C_1X_2N_{21} + C_2X_2N_{21}{}^{D} \underset{koff_{p1}}{\overset{kon_{p1}}{\rightleftharpoons}} C_1X_2N_{21}{-}C_2X_2N_{21}{}^{D} \tag{136}$$

$$C_2X_1N_{12} + C_1X_1N_{12}{}^{D} \underset{koff_{p2}}{\overset{kon_{p2}}{\rightleftharpoons}} C_2X_1N_{12}{-}C_1X_1N_{12}{}^{D} \tag{137}$$

$$C_2X_1N_{12} + C_1X_2N_{22}{}^{D} \underset{koff_{p2}}{\overset{kon_{p2}}{\rightleftharpoons}} C_2X_1N_{12}{-}C_1X_2N_{22}{}^{D} \tag{138}$$

$$C_2X_1N_{12} + C_2X_1N_{12}{}^{D} \underset{koff_{p2}}{\overset{kon_{p2}}{\rightleftharpoons}} C_2X_1N_{12}{-}C_2X_1N_{12}{}^{D} \tag{139}$$

$$C_2X_1N_{12} + C_2X_2N_{22}{}^{D} \underset{koff_{p2}}{\overset{kon_{p2}}{\rightleftharpoons}} C_2X_1N_{12}{-}C_2X_2N_{22}{}^{D} \tag{140}$$

$$C_2X_2N_{22} + C_1X_1N_{12}{}^{D} \underset{koff_{p2}}{\overset{kon_{p2}}{\rightleftharpoons}} C_2X_2N_{22}{-}C_1X_1N_{12}{}^{D} \tag{141}$$

$$C_2X_2N_{22} + C_1X_2N_{22}{}^{D} \underset{koff_{p2}}{\overset{kon_{p2}}{\rightleftharpoons}} C_2X_2N_{22}{-}C_1X_2N_{22}{}^{D} \tag{142}$$

$$C_2X_2N_{22} + C_2X_1N_{12}{}^{D} \underset{koff_{p2}}{\overset{kon_{p2}}{\rightleftharpoons}} C_2X_2N_{22}{-}C_2X_1N_{12}{}^{D} \tag{143}$$

$$C_2X_2N_{22} + C_2X_2N_{22}{}^{D} \underset{koff_{p2}}{\overset{kon_{p2}}{\rightleftharpoons}} C_2X_2N_{22}{-}C_2X_2N_{22}{}^{D} \tag{144}$$

$$C_1X_1N_{11}{-}C_1X_1N_{11}{}^{D} \xrightarrow{kcat_1} 2\,C_1X_1N_{11} \tag{145}$$

$$C_1X_1N_{11}{-}C_1X_2N_{21}{}^{D} \xrightarrow{kcat_1} C_1X_1N_{11} + C_1X_2N_{21} \tag{146}$$

$$C_1X_1N_{11}{-}C_2X_1N_{11}{}^{D} \xrightarrow{kcat_1} C_1X_1N_{11} + C_2X_1N_{11} \tag{147}$$

$$C_1X_1N_{11}{-}C_2X_2N_{21}{}^{D} \xrightarrow{kcat_1} C_1X_1N_{11} + C_2X_2N_{21} \tag{148}$$

$$C_1X_2N_{21}{-}C_1X_1N_{11}{}^{D} \xrightarrow{kcat_1} C_1X_2N_{21} + C_1X_1N_{11} \tag{149}$$

$$C_1X_2N_{21}{-}C_1X_2N_{21}{}^{D} \xrightarrow{kcat_1} 2\,C_1X_2N_{21} \tag{150}$$

$$\mathrm{C_1X_2N_{21}{-}C_2X_1N_{11}}^{\mathrm{D}} \xrightarrow{\mathrm{kcat_1}} \mathrm{C_1X_2N_{21}} + \mathrm{C_2X_1N_{11}} \tag{151}$$

$$\mathrm{C_1X_2N_{21}{-}C_2X_2N_{21}}^{\mathrm{D}} \xrightarrow{\mathrm{kcat_1}} \mathrm{C_1X_2N_{21}} + \mathrm{C_2X_2N_{21}} \tag{152}$$

$$\mathrm{C_2X_1N_{12}{-}C_1X_1N_{12}}^{\mathrm{D}} \xrightarrow{\mathrm{kcat_2}} \mathrm{C_2X_1N_{12}} + \mathrm{C_1X_1N_{12}} \tag{153}$$

$$\mathrm{C_2X_1N_{12}{-}C_1X_2N_{22}}^{\mathrm{D}} \xrightarrow{\mathrm{kcat_2}} \mathrm{C_2X_1N_{12}} + \mathrm{C_1X_2N_{22}} \tag{154}$$

$$\mathrm{C_2X_1N_{12}{-}C_2X_1N_{12}}^{\mathrm{D}} \xrightarrow{\mathrm{kcat_2}} 2\,\mathrm{C_2X_1N_{12}} \tag{155}$$

$$\mathrm{C_2X_1N_{12}{-}C_2X_2N_{22}}^{\mathrm{D}} \xrightarrow{\mathrm{kcat_2}} \mathrm{C_2X_1N_{12}} + \mathrm{C_2X_2N_{22}} \tag{156}$$

$$\mathrm{C_2X_2N_{22}{-}C_1X_1N_{12}}^{\mathrm{D}} \xrightarrow{\mathrm{kcat_2}} \mathrm{C_2X_2N_{22}} + \mathrm{C_1X_1N_{12}} \tag{157}$$

$$\mathrm{C_2X_2N_{22}{-}C_1X_2N_{22}}^{\mathrm{D}} \xrightarrow{\mathrm{kcat_2}} \mathrm{C_2X_2N_{22}} + \mathrm{C_1X_2N_{22}} \tag{158}$$

$$\mathrm{C_2X_2N_{22}{-}C_2X_1N_{12}}^{\mathrm{D}} \xrightarrow{\mathrm{kcat_2}} \mathrm{C_2X_2N_{22}} + \mathrm{C_2X_1N_{12}} \tag{159}$$

$$\mathrm{C_2X_2N_{22}{-}C_2X_2N_{22}}^{\mathrm{D}} \xrightarrow{\mathrm{kcat_2}} 2\,\mathrm{C_2X_2N_{22}} \tag{160}$$

$$\mathrm{C_1X_1N_{11}} + \mathrm{C_2} \underset{\mathrm{koff}_{p1}}{\overset{\mathrm{kon}_{p1}}{\rightleftharpoons}} \mathrm{C_1X_1N_{11}{-}C_2} \tag{161}$$

$$\mathrm{C_1X_2N_{21}} + \mathrm{C_2} \underset{\mathrm{koff}_{p1}}{\overset{\mathrm{kon}_{p1}}{\rightleftharpoons}} \mathrm{C_1X_2N_{21}{-}C_2} \tag{162}$$

$$\mathrm{C_2X_1N_{12}} + \mathrm{C_1} \underset{\mathrm{koff}_{p2}}{\overset{\mathrm{kon}_{p2}}{\rightleftharpoons}} \mathrm{C_2X_1N_{12}{-}C_1} \tag{163}$$

$$\mathrm{C_2X_2N_{22}} + \mathrm{C_1} \underset{\mathrm{koff}_{p2}}{\overset{\mathrm{kon}_{p2}}{\rightleftharpoons}} \mathrm{C_2X_2N_{22}{-}C_1} \tag{164}$$

$$\mathrm{C_1X_1N_{11}{-}C_2} \xrightarrow{\mathrm{kcat_1}} \mathrm{C_1X_1N_{11}} + \mathrm{C_2}^{\mathrm{E}} \tag{165}$$

$$\mathrm{C_1X_2N_{21}{-}C_2} \xrightarrow{\mathrm{kcat_1}} \mathrm{C_1X_2N_{21}} + \mathrm{C_2}^{\mathrm{E}} \tag{166}$$

$$\mathrm{C_2X_1N_{12}{-}C_1} \xrightarrow{\mathrm{kcat_2}} \mathrm{C_2X_1N_{12}} + \mathrm{C_1}^{\mathrm{E}} \tag{167}$$

$$\mathrm{C_2X_2N_{22}{-}C_1} \xrightarrow{\mathrm{kcat_2}} \mathrm{C_2X_2N_{22}} + \mathrm{C_1}^{\mathrm{E}} \tag{168}$$

$$\mathrm{C_2}^{\mathrm{E}} + \mathrm{X_1N_{11}}^{\mathrm{D}} \underset{\mathrm{koff_2}}{\overset{\mathrm{kon_2}}{\rightleftharpoons}} \mathrm{C_2}^{\mathrm{E}}\mathrm{X_1N_{11}}^{\mathrm{D}} \tag{169}$$

$$\mathrm{C_2}^{\mathrm{E}} + \mathrm{X_1N_{12}}^{\mathrm{D}} \underset{\mathrm{koff_2}}{\overset{\mathrm{kon_2}}{\rightleftharpoons}} \mathrm{C_2}^{\mathrm{E}}\mathrm{X_1N_{12}}^{\mathrm{D}} \tag{170}$$

$$\mathrm{C_2}^{\mathrm{E}} + \mathrm{X_2N_{21}}^{\mathrm{D}} \underset{\mathrm{koff_2}}{\overset{\mathrm{kon_2}}{\rightleftharpoons}} \mathrm{C_2}^{\mathrm{E}}\mathrm{X_2N_{21}}^{\mathrm{D}} \tag{171}$$

$$\mathrm{C_2}^{\mathrm{E}} + \mathrm{X_2N_{22}}^{\mathrm{D}} \underset{\mathrm{koff_2}}{\overset{\mathrm{kon_2}}{\rightleftharpoons}} \mathrm{C_2}^{\mathrm{E}}\mathrm{X_2N_{22}}^{\mathrm{D}} \tag{172}$$

$$\mathrm{C_1}^{\mathrm{E}} + \mathrm{X_1N_{11}}^{\mathrm{D}} \underset{\mathrm{koff_2}}{\overset{\mathrm{kon_2}}{\rightleftharpoons}} \mathrm{C_1}^{\mathrm{E}}\mathrm{X_1N_{11}}^{\mathrm{D}} \tag{173}$$

$$\mathrm{C_1}^{\mathrm{E}} + \mathrm{X_1N_{12}}^{\mathrm{D}} \underset{\mathrm{koff_2}}{\overset{\mathrm{kon_2}}{\rightleftharpoons}} \mathrm{C_1}^{\mathrm{E}}\mathrm{X_1N_{12}}^{\mathrm{D}} \tag{174}$$

$$\mathrm{C_1}^{\mathrm{E}} + \mathrm{X_2N_{21}}^{\mathrm{D}} \underset{\mathrm{koff_2}}{\overset{\mathrm{kon_2}}{\rightleftharpoons}} \mathrm{C_1}^{\mathrm{E}}\mathrm{X_2N_{21}}^{\mathrm{D}} \tag{175}$$

$$C_1{}^E + X_2N_{22}{}^D \underset{koff_2}{\overset{kon_2}{\rightleftharpoons}} C_1{}^EX_2N_{22}{}^D \quad (176)$$

$$C_2{}^E + X_1N_{11} \underset{koff_2}{\overset{kon_2}{\rightleftharpoons}} C_2{}^EX_1N_{11} \quad (177)$$

$$C_2{}^E + X_2N_{21} \underset{koff_2}{\overset{kon_2}{\rightleftharpoons}} C_2{}^EX_2N_{21} \quad (178)$$

$$C_2{}^E + X_1N_{12} \underset{koff_2}{\overset{kon_2}{\rightleftharpoons}} C_2{}^EX_1N_{12} \quad (179)$$

$$C_2{}^E + X_2N_{22} \underset{koff_2}{\overset{kon_2}{\rightleftharpoons}} C_2{}^EX_2N_{22} \quad (180)$$

$$C_1{}^E + X_1N_{11} \underset{koff_2}{\overset{kon_2}{\rightleftharpoons}} C_1{}^EX_1N_{11} \quad (181)$$

$$C_1{}^E + X_2N_{21} \underset{koff_2}{\overset{kon_2}{\rightleftharpoons}} C_1{}^EX_2N_{21} \quad (182)$$

$$C_1{}^E + X_1N_{12} \underset{koff_2}{\overset{kon_2}{\rightleftharpoons}} C_1{}^EX_1N_{12} \quad (183)$$

$$C_1{}^E + X_2N_{22} \underset{koff_2}{\overset{kon_2}{\rightleftharpoons}} C_1{}^EX_2N_{22} \quad (184)$$

$$C_1X_1N_{11}{}^D + C_2{}^EX_1N_{11}{}^D \underset{koff_{p1}}{\overset{kon_{p1}}{\rightleftharpoons}} C_1X_1N_{11}{}^D{-}C_2{}^EX_1N_{11}{}^D \quad (185)$$

$$C_1X_1N_{11}{}^D + C_2{}^EX_2N_{21}{}^D \underset{koff_{p1}}{\overset{kon_{p1}}{\rightleftharpoons}} C_1X_1N_{11}{}^D{-}C_2{}^EX_2N_{21}{}^D \quad (186)$$

$$C_1X_1N_{11}{}^D + C_1{}^EX_1N_{11}{}^D \underset{koff_{p1}}{\overset{kon_{p1}}{\rightleftharpoons}} C_1X_1N_{11}{}^D{-}C_1{}^EX_1N_{11}{}^D \quad (187)$$

$$C_1X_1N_{11}{}^D + C_1{}^EX_2N_{21}{}^D \underset{koff_{p1}}{\overset{kon_{p1}}{\rightleftharpoons}} C_1X_1N_{11}{}^D{-}C_1{}^EX_2N_{21}{}^D \quad (188)$$

$$C_1X_2N_{21}{}^D + C_2{}^EX_1N_{11}{}^D \underset{koff_{p1}}{\overset{kon_{p1}}{\rightleftharpoons}} C_1X_2N_{21}{}^D{-}C_2{}^EX_1N_{11}{}^D \quad (189)$$

$$C_1X_2N_{21}{}^D + C_2{}^EX_2N_{21}{}^D \underset{koff_{p1}}{\overset{kon_{p1}}{\rightleftharpoons}} C_1X_2N_{21}{}^D{-}C_2{}^EX_2N_{21}{}^D \quad (190)$$

$$C_1X_2N_{21}{}^D + C_1{}^EX_1N_{11}{}^D \underset{koff_{p1}}{\overset{kon_{p1}}{\rightleftharpoons}} C_1X_2N_{21}{}^D{-}C_1{}^EX_1N_{11}{}^D \quad (191)$$

$$C_1X_2N_{21}{}^D + C_1{}^EX_2N_{21}{}^D \underset{koff_{p1}}{\overset{kon_{p1}}{\rightleftharpoons}} C_1X_2N_{21}{}^D{-}C_1{}^EX_2N_{21}{}^D \quad (192)$$

$$C_2X_1N_{12}{}^D + C_2{}^EX_1N_{12}{}^D \underset{koff_{p2}}{\overset{kon_{p2}}{\rightleftharpoons}} C_2X_1N_{12}{}^D{-}C_2{}^EX_1N_{12}{}^D \quad (193)$$

$$C_2X_1N_{12}{}^D + C_2{}^EX_2N_{22}{}^D \underset{koff_{p2}}{\overset{kon_{p2}}{\rightleftharpoons}} C_2X_1N_{12}{}^D{-}C_2{}^EX_2N_{22}{}^D \quad (194)$$

$$C_2X_1N_{12}{}^D + C_1{}^EX_1N_{12}{}^D \underset{koff_{p2}}{\overset{kon_{p2}}{\rightleftharpoons}} C_2X_1N_{12}{}^D{-}C_1{}^EX_1N_{12}{}^D \quad (195)$$

$$C_2X_1N_{12}{}^D + C_1{}^EX_2N_{22}{}^D \underset{koff_{p2}}{\overset{kon_{p2}}{\rightleftharpoons}} C_2X_1N_{12}{}^D{-}C_1{}^EX_2N_{22}{}^D \quad (196)$$

$$C_2X_2N_{22}{}^D + C_2{}^EX_1N_{12}{}^D \underset{koff_{p2}}{\overset{kon_{p2}}{\rightleftharpoons}} C_2X_2N_{22}{}^D{-}C_2{}^EX_1N_{12}{}^D \quad (197)$$

$$C_2X_2N_{22}{}^D + C_2{}^EX_2N_{22}{}^D \underset{koff_{p2}}{\overset{kon_{p2}}{\rightleftharpoons}} C_2X_2N_{22}{}^D{-}C_2{}^EX_2N_{22}{}^D \quad (198)$$

$$C_2X_2N_{22}{}^D + C_1{}^EX_1N_{12}{}^D \underset{koff_{p2}}{\overset{kon_{p2}}{\rightleftharpoons}} C_2X_2N_{22}{}^D{-}C_1{}^EX_1N_{12}{}^D \quad (199)$$

$$C_2X_2N_{22}{}^D + C_1{}^EX_2N_{22}{}^D \underset{koff_{p2}}{\overset{kon_{p2}}{\rightleftharpoons}} C_2X_2N_{22}{}^D{-}C_1{}^EX_2N_{22}{}^D \quad (200)$$

$$C_1X_1N_{11}{}^{D}{-}C_2{}^{E}X_1N_{11}{}^{D} \xrightarrow{kcat_1} C_1X_1N_{11}{}^{D} + C_2{}^{E}X_1N_{11} \tag{201}$$

$$C_1X_1N_{11}{}^{D}{-}C_2{}^{E}X_2N_{21}{}^{D} \xrightarrow{kcat_1} C_1X_1N_{11}{}^{D} + C_2{}^{E}X_2N_{21} \tag{202}$$

$$C_1X_1N_{11}{}^{D}{-}C_1{}^{E}X_1N_{11}{}^{D} \xrightarrow{kcat_1} C_1X_1N_{11}{}^{D} + C_1{}^{E}X_1N_{11} \tag{203}$$

$$C_1X_1N_{11}{}^{D}{-}C_1{}^{E}X_2N_{21}{}^{D} \xrightarrow{kcat_1} C_1X_1N_{11}{}^{D} + C_1{}^{E}X_2N_{21} \tag{204}$$

$$C_1X_2N_{21}{}^{D}{-}C_2{}^{E}X_1N_{11}{}^{D} \xrightarrow{kcat_1} C_1X_2N_{21}{}^{D} + C_2{}^{E}X_1N_{11} \tag{205}$$

$$C_1X_2N_{21}{}^{D}{-}C_2{}^{E}X_2N_{21}{}^{D} \xrightarrow{kcat_1} C_1X_2N_{21}{}^{D} + C_2{}^{E}X_2N_{21} \tag{206}$$

$$C_1X_2N_{21}{}^{D}{-}C_1{}^{E}X_1N_{11}{}^{D} \xrightarrow{kcat_1} C_1X_2N_{21}{}^{D} + C_1{}^{E}X_1N_{11} \tag{207}$$

$$C_1X_2N_{21}{}^{D}{-}C_1{}^{E}X_2N_{21}{}^{D} \xrightarrow{kcat_1} C_1X_2N_{21}{}^{D} + C_1{}^{E}X_2N_{21} \tag{208}$$

$$C_2X_1N_{12}{}^{D}{-}C_2{}^{E}X_1N_{12}{}^{D} \xrightarrow{kcat_2} C_2X_1N_{12}{}^{D} + C_2{}^{E}X_1N_{12} \tag{209}$$

$$C_2X_1N_{12}{}^{D}{-}C_2{}^{E}X_2N_{22}{}^{D} \xrightarrow{kcat_2} C_2X_1N_{12}{}^{D} + C_2{}^{E}X_2N_{22} \tag{210}$$

$$C_2X_1N_{12}{}^{D}{-}C_1{}^{E}X_1N_{12}{}^{D} \xrightarrow{kcat_2} C_2X_1N_{12}{}^{D} + C_1{}^{E}X_1N_{12} \tag{211}$$

$$C_2X_1N_{12}{}^{D}{-}C_1{}^{E}X_2N_{22}{}^{D} \xrightarrow{kcat_2} C_2X_1N_{12}{}^{D} + C_1{}^{E}X_2N_{22} \tag{212}$$

$$C_2X_2N_{22}{}^{D}{-}C_2{}^{E}X_1N_{12}{}^{D} \xrightarrow{kcat_2} C_2X_2N_{22}{}^{D} + C_2{}^{E}X_1N_{12} \tag{213}$$

$$C_2X_2N_{22}{}^{D}{-}C_2{}^{E}X_2N_{22}{}^{D} \xrightarrow{kcat_2} C_2X_2N_{22}{}^{D} + C_2{}^{E}X_2N_{22} \tag{214}$$

$$C_2X_2N_{22}{}^{D}{-}C_1{}^{E}X_1N_{12}{}^{D} \xrightarrow{kcat_2} C_2X_2N_{22}{}^{D} + C_1{}^{E}X_1N_{12} \tag{215}$$

$$C_2X_2N_{22}{}^{D}{-}C_1{}^{E}X_2N_{22}{}^{D} \xrightarrow{kcat_2} C_2X_2N_{22}{}^{D} + C_1{}^{E}X_2N_{22} \tag{216}$$

$$C_1X_1N_{11} + C_2{}^{E}X_1N_{11}{}^{D} \underset{koff_{p1}}{\overset{kon_{p1}}{\rightleftharpoons}} C_1X_1N_{11}{-}C_2{}^{E}X_1N_{11}{}^{D} \tag{217}$$

$$C_1X_1N_{11} + C_2{}^{E}X_2N_{21}{}^{D} \underset{koff_{p1}}{\overset{kon_{p1}}{\rightleftharpoons}} C_1X_1N_{11}{-}C_2{}^{E}X_2N_{21}{}^{D} \tag{218}$$

$$C_1X_1N_{11} + C_1{}^{E}X_1N_{11}{}^{D} \underset{koff_{p1}}{\overset{kon_{p1}}{\rightleftharpoons}} C_1X_1N_{11}{-}C_1{}^{E}X_1N_{11}{}^{D} \tag{219}$$

$$C_1X_1N_{11} + C_1{}^{E}X_2N_{21}{}^{D} \underset{koff_{p1}}{\overset{kon_{p1}}{\rightleftharpoons}} C_1X_1N_{11}{-}C_1{}^{E}X_2N_{21}{}^{D} \tag{220}$$

$$C_1X_2N_{21} + C_2{}^{E}X_1N_{11}{}^{D} \underset{koff_{p1}}{\overset{kon_{p1}}{\rightleftharpoons}} C_1X_2N_{21}{-}C_2{}^{E}X_1N_{11}{}^{D} \tag{221}$$

$$C_1X_2N_{21} + C_2{}^{E}X_2N_{21}{}^{D} \underset{koff_{p1}}{\overset{kon_{p1}}{\rightleftharpoons}} C_1X_2N_{21}{-}C_2{}^{E}X_2N_{21}{}^{D} \tag{222}$$

$$C_1X_2N_{21} + C_1{}^{E}X_1N_{11}{}^{D} \underset{koff_{p1}}{\overset{kon_{p1}}{\rightleftharpoons}} C_1X_2N_{21}{-}C_1{}^{E}X_1N_{11}{}^{D} \tag{223}$$

$$C_1X_2N_{21} + C_1{}^{E}X_2N_{21}{}^{D} \underset{koff_{p1}}{\overset{kon_{p1}}{\rightleftharpoons}} C_1X_2N_{21}{-}C_1{}^{E}X_2N_{21}{}^{D} \tag{224}$$

$$C_2X_1N_{12} + C_2{}^{E}X_1N_{12}{}^{D} \underset{koff_{p2}}{\overset{kon_{p2}}{\rightleftharpoons}} C_2X_1N_{12}{-}C_2{}^{E}X_1N_{12}{}^{D} \tag{225}$$

$$C_2X_1N_{12} + C_2{}^EX_2N_{22}{}^D \underset{koff_{p2}}{\overset{kon_{p2}}{\rightleftharpoons}} C_2X_1N_{12}{-}C_2{}^EX_2N_{22}{}^D \tag{226}$$

$$C_2X_1N_{12} + C_1{}^EX_1N_{12}{}^D \underset{koff_{p2}}{\overset{kon_{p2}}{\rightleftharpoons}} C_2X_1N_{12}{-}C_1{}^EX_1N_{12}{}^D \tag{227}$$

$$C_2X_1N_{12} + C_1{}^EX_2N_{22}{}^D \underset{koff_{p2}}{\overset{kon_{p2}}{\rightleftharpoons}} C_2X_1N_{12}{-}C_1{}^EX_2N_{22}{}^D \tag{228}$$

$$C_2X_2N_{22} + C_2{}^EX_1N_{12}{}^D \underset{koff_{p2}}{\overset{kon_{p2}}{\rightleftharpoons}} C_2X_2N_{22}{-}C_2{}^EX_1N_{12}{}^D \tag{229}$$

$$C_2X_2N_{22} + C_2{}^EX_2N_{22}{}^D \underset{koff_{p2}}{\overset{kon_{p2}}{\rightleftharpoons}} C_2X_2N_{22}{-}C_2{}^EX_2N_{22}{}^D \tag{230}$$

$$C_2X_2N_{22} + C_1{}^EX_1N_{12}{}^D \underset{koff_{p2}}{\overset{kon_{p2}}{\rightleftharpoons}} C_2X_2N_{22}{-}C_1{}^EX_1N_{12}{}^D \tag{231}$$

$$C_2X_2N_{22} + C_1{}^EX_2N_{22}{}^D \underset{koff_{p2}}{\overset{kon_{p2}}{\rightleftharpoons}} C_2X_2N_{22}{-}C_1{}^EX_2N_{22}{}^D \tag{232}$$

$$C_1X_1N_{11}{-}C_2{}^EX_1N_{11}{}^D \xrightarrow{kcat_1} C_1X_1N_{11} + C_2{}^EX_1N_{11} \tag{233}$$

$$C_1X_1N_{11}{-}C_2{}^EX_2N_{21}{}^D \xrightarrow{kcat_1} C_1X_1N_{11} + C_2{}^EX_2N_{21} \tag{234}$$

$$C_1X_1N_{11}{-}C_1{}^EX_1N_{11}{}^D \xrightarrow{kcat_1} C_1X_1N_{11} + C_1{}^EX_1N_{11} \tag{235}$$

$$C_1X_1N_{11}{-}C_1{}^EX_2N_{21}{}^D \xrightarrow{kcat_1} C_1X_1N_{11} + C_1{}^EX_2N_{21} \tag{236}$$

$$C_1X_2N_{21}{-}C_2{}^EX_1N_{11}{}^D \xrightarrow{kcat_1} C_1X_2N_{21} + C_2{}^EX_1N_{11} \tag{237}$$

$$C_1X_2N_{21}{-}C_2{}^EX_2N_{21}{}^D \xrightarrow{kcat_1} C_1X_2N_{21} + C_2{}^EX_2N_{21} \tag{238}$$

$$C_1X_2N_{21}{-}C_1{}^EX_1N_{11}{}^D \xrightarrow{kcat_1} C_1X_2N_{21} + C_1{}^EX_1N_{11} \tag{239}$$

$$C_1X_2N_{21}{-}C_1{}^EX_2N_{21}{}^D \xrightarrow{kcat_1} C_1X_2N_{21} + C_1{}^EX_2N_{21} \tag{240}$$

$$C_2X_1N_{12}{-}C_2{}^EX_1N_{12}{}^D \xrightarrow{kcat_2} C_2X_1N_{12} + C_2{}^EX_1N_{12} \tag{241}$$

$$C_2X_1N_{12}{-}C_2{}^EX_2N_{22}{}^D \xrightarrow{kcat_2} C_2X_1N_{12} + C_2{}^EX_2N_{22} \tag{242}$$

$$C_2X_1N_{12}{-}C_1{}^EX_1N_{12}{}^D \xrightarrow{kcat_2} C_2X_1N_{12} + C_1{}^EX_1N_{12} \tag{243}$$

$$C_2X_1N_{12}{-}C_1{}^EX_2N_{22}{}^D \xrightarrow{kcat_2} C_2X_1N_{12} + C_1{}^EX_2N_{22} \tag{244}$$

$$C_2X_2N_{22}{-}C_2{}^EX_1N_{12}{}^D \xrightarrow{kcat_2} C_2X_2N_{22} + C_2{}^EX_1N_{12} \tag{245}$$

$$C_2X_2N_{22}{-}C_2{}^EX_2N_{22}{}^D \xrightarrow{kcat_2} C_2X_2N_{22} + C_2{}^EX_2N_{22} \tag{246}$$

$$C_2X_2N_{22}{-}C_1{}^EX_1N_{12}{}^D \xrightarrow{kcat_2} C_2X_2N_{22} + C_1{}^EX_1N_{12} \tag{247}$$

$$C_2X_2N_{22}{-}C_1{}^EX_2N_{22}{}^D \xrightarrow{kcat_2} C_2X_2N_{22} + C_1{}^EX_2N_{22} \tag{248}$$

$$\varnothing \xrightarrow{syn_{X1}\,/(avgd \cdot cell)} X_1 \tag{249}$$

$$\varnothing \xrightarrow{syn_{X2}\,/(avgd \cdot cell)} X_2 \tag{250}$$

$$\varnothing \xrightarrow{\mathrm{syn_{N11}}\ /(\mathrm{avgd}\cdot\mathrm{cell})} \mathrm{N_{11}}^{\mathrm{D}} \tag{251}$$

$$\varnothing \xrightarrow{\mathrm{syn_{N12}}\ /(\mathrm{avgd}\cdot\mathrm{cell})} \mathrm{N_{12}}^{\mathrm{D}} \tag{252}$$

$$\varnothing \xrightarrow{\mathrm{syn_{N21}}\ /(\mathrm{avgd}\cdot\mathrm{cell})} \mathrm{N_{21}}^{\mathrm{D}} \tag{253}$$

$$\varnothing \xrightarrow{\mathrm{syn_{N22}}\ /(\mathrm{avgd}\cdot\mathrm{cell})} \mathrm{N_{22}}^{\mathrm{D}} \tag{254}$$

$$\varnothing \xrightarrow{\mathrm{syn{-}C_1}\ /(\mathrm{avgd}\cdot\mathrm{cell})} \mathrm{C_1} \tag{255}$$

$$\varnothing \xrightarrow{\mathrm{syn{-}C_2}\ /(\mathrm{avgd}\cdot\mathrm{cell})} \mathrm{C_2} \tag{256}$$

$$\mathrm{X_1N_{11}}^{\mathrm{D}} \xrightarrow{\mathrm{deg_{DHFR}}} \varnothing \tag{257}$$

$$\mathrm{X_1N_{12}}^{\mathrm{D}} \xrightarrow{\mathrm{deg_{DHFR}}} \varnothing \tag{258}$$

$$\mathrm{X_2N_{21}}^{\mathrm{D}} \xrightarrow{\mathrm{deg_{DHFR}}} \varnothing \tag{259}$$

$$\mathrm{X_2N_{22}}^{\mathrm{D}} \xrightarrow{\mathrm{deg_{DHFR}}} \varnothing \tag{260}$$

$$\mathrm{C_1X_1N_{11}}^{\mathrm{D}} \xrightarrow{\mathrm{deg_{DHFR}}} \varnothing \tag{261}$$

$$\mathrm{C_1X_1N_{12}}^{\mathrm{D}} \xrightarrow{\mathrm{deg_{DHFR}}} \varnothing \tag{262}$$

$$\mathrm{C_1X_2N_{21}}^{\mathrm{D}} \xrightarrow{\mathrm{deg_{DHFR}}} \varnothing \tag{263}$$

$$\mathrm{C_1X_2N_{22}}^{\mathrm{D}} \xrightarrow{\mathrm{deg_{DHFR}}} \varnothing \tag{264}$$

$$\mathrm{C_2X_1N_{11}}^{\mathrm{D}} \xrightarrow{\mathrm{deg_{DHFR}}} \varnothing \tag{265}$$

$$\mathrm{C_2X_1N_{12}}^{\mathrm{D}} \xrightarrow{\mathrm{deg_{DHFR}}} \varnothing \tag{266}$$

$$\mathrm{C_2X_2N_{21}}^{\mathrm{D}} \xrightarrow{\mathrm{deg_{DHFR}}} \varnothing \tag{267}$$

$$\mathrm{C_2X_2N_{22}}^{\mathrm{D}} \xrightarrow{\mathrm{deg_{DHFR}}} \varnothing \tag{268}$$

$$\mathrm{N_{11}} \xrightarrow{\mathrm{deg_{reg}}} \varnothing \tag{269}$$

$$\mathrm{N_{21}} \xrightarrow{\mathrm{deg_{reg}}} \varnothing \tag{270}$$

$$\mathrm{N_{12}} \xrightarrow{\mathrm{deg_{reg}}} \varnothing \tag{271}$$

$$\mathrm{N_{22}} \xrightarrow{\mathrm{deg_{reg}}} \varnothing \tag{272}$$

$$\mathrm{X_1N_{11}} \xrightarrow{\mathrm{deg_{reg}}} \varnothing \tag{273}$$

$$\mathrm{X_2N_{21}} \xrightarrow{\mathrm{deg_{reg}}} \varnothing \tag{274}$$

$$\mathrm{X_1N_{12}} \xrightarrow{\mathrm{deg_{reg}}} \varnothing \tag{275}$$

$$X_2N_{22} \xrightarrow{\text{deg}_{\text{reg}}} \varnothing \tag{276}$$

$$C_1X_1N_{11} \xrightarrow{\text{deg}_{\text{reg}}} \varnothing \tag{277}$$

$$C_1X_2N_{21} \xrightarrow{\text{deg}_{\text{reg}}} \varnothing \tag{278}$$

$$C_2X_1N_{11} \xrightarrow{\text{deg}_{\text{reg}}} \varnothing \tag{279}$$

$$C_2X_2N_{21} \xrightarrow{\text{deg}_{\text{reg}}} \varnothing \tag{280}$$

$$C_1X_1N_{12} \xrightarrow{\text{deg}_{\text{reg}}} \varnothing \tag{281}$$

$$C_1X_2N_{22} \xrightarrow{\text{deg}_{\text{reg}}} \varnothing \tag{282}$$

$$C_2X_1N_{12} \xrightarrow{\text{deg}_{\text{reg}}} \varnothing \tag{283}$$

$$C_2X_2N_{22} \xrightarrow{\text{deg}_{\text{reg}}} \varnothing \tag{284}$$

$${C_2}^E \xrightarrow{\text{deg}_{\text{reg}}} \varnothing \tag{285}$$

$${C_1}^E \xrightarrow{\text{deg}_{\text{reg}}} \varnothing \tag{286}$$

$${C_2}^E X_1 {N_{11}}^D \xrightarrow{\text{deg}_{\text{DHFR}}} \varnothing \tag{287}$$

$${C_2}^E X_1 {N_{12}}^D \xrightarrow{\text{deg}_{\text{DHFR}}} \varnothing \tag{288}$$

$${C_2}^E X_2 {N_{21}}^D \xrightarrow{\text{deg}_{\text{DHFR}}} \varnothing \tag{289}$$

$${C_2}^E X_2 {N_{22}}^D \xrightarrow{\text{deg}_{\text{DHFR}}} \varnothing \tag{290}$$

$${C_1}^E X_1 {N_{11}}^D \xrightarrow{\text{deg}_{\text{DHFR}}} \varnothing \tag{291}$$

$${C_1}^E X_1 {N_{12}}^D \xrightarrow{\text{deg}_{\text{DHFR}}} \varnothing \tag{292}$$

$${C_1}^E X_2 {N_{21}}^D \xrightarrow{\text{deg}_{\text{DHFR}}} \varnothing \tag{293}$$

$${C_1}^E X_2 {N_{22}}^D \xrightarrow{\text{deg}_{\text{DHFR}}} \varnothing \tag{294}$$

$${C_2}^E X_1 N_{11} \xrightarrow{\text{deg}_{\text{reg}}} \varnothing \tag{295}$$

$${C_2}^E X_2 N_{21} \xrightarrow{\text{deg}_{\text{reg}}} \varnothing \tag{296}$$

$${C_1}^E X_1 N_{11} \xrightarrow{\text{deg}_{\text{reg}}} \varnothing \tag{297}$$

$${C_1}^E X_2 N_{21} \xrightarrow{\text{deg}_{\text{reg}}} \varnothing \tag{298}$$

$${C_2}^E X_1 N_{12} \xrightarrow{\text{deg}_{\text{reg}}} \varnothing \tag{299}$$

$${C_2}^E X_2 N_{22} \xrightarrow{\text{deg}_{\text{reg}}} \varnothing \tag{300}$$

$${C_1}^E X_1 N_{12} \xrightarrow{\text{deg}_{\text{reg}}} \varnothing \tag{301}$$

$${C_1}^E X_2 N_{22} \xrightarrow{\text{deg}_{\text{reg}}} \varnothing \tag{302}$$

$$X_1 \xrightarrow{\text{deg}_{\text{reg}}} \varnothing \tag{303}$$

$$X_2 \xrightarrow{\text{deg}_{\text{reg}}} \varnothing \tag{304}$$

$${N_{11}}^D \xrightarrow{\text{deg}_{\text{DHFR}}} \varnothing \tag{305}$$

$${N_{12}}^D \xrightarrow{\text{deg}_{\text{DHFR}}} \varnothing \tag{306}$$

$${N_{21}}^D \xrightarrow{\text{deg}_{\text{DHFR}}} \varnothing \tag{307}$$

$${N_{22}}^D \xrightarrow{\text{deg}_{\text{DHFR}}} \varnothing \tag{308}$$

$$C_1 \xrightarrow{\text{deg}_{\text{reg}}} \varnothing \tag{309}$$

$$C_2 \xrightarrow{\text{deg}_{\text{reg}}} \varnothing \tag{310}$$

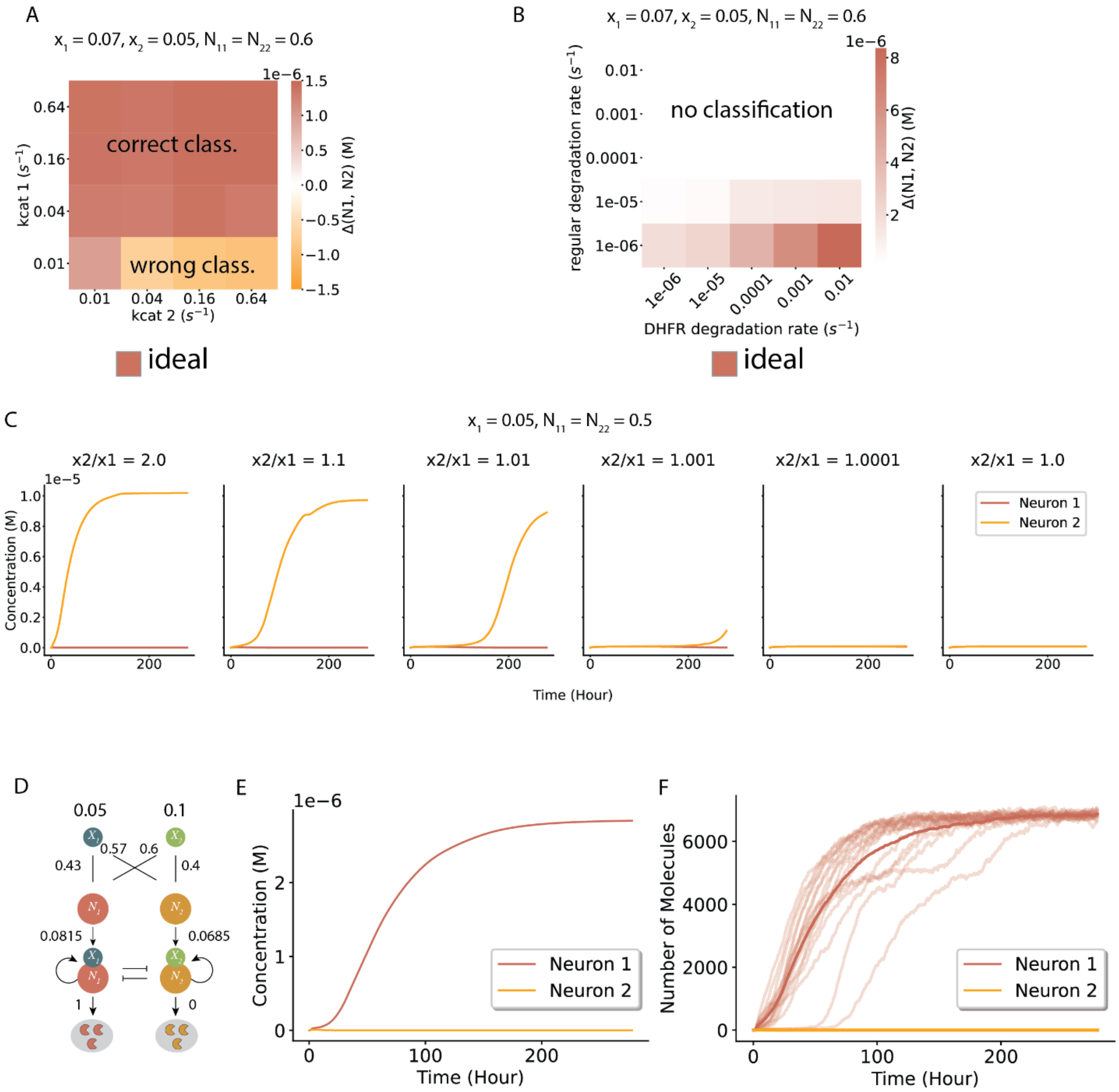

A
$x_1 = 0.07, x_2 = 0.05, N_{11} = N_{22} = 0.6$
correct class.
wrong class.
kcat 1 ($s^{-1}$)
kcat 2 ($s^{-1}$)
Δ(N1, N2) (M)
ideal
B
$x_1 = 0.07, x_2 = 0.05, N_{11} = N_{22} = 0.6$
no classification
regular degradation rate ($s^{-1}$)
DHFR degradation rate ($s^{-1}$)
Δ(N1, N2) (M)
ideal
C
$x_1 = 0.05, N_{11} = N_{22} = 0.5$
x2/x1 = 2.0
x2/x1 = 1.1
x2/x1 = 1.01
x2/x1 = 1.001
x2/x1 = 1.0001
x2/x1 = 1.0
Concentration (M)
Time (Hour)
Neuron 1
Neuron 2
D
0.05
0.1
0.57
0.6
0.43
0.4
0.0815
0.0685
1
0
E
Concentration (M)
Time (Hour)
Neuron 1
Neuron 2
F
Number of Molecules
Time (Hour)
Neuron 1
Neuron 2

**Figure S1. Simulations of the winner-take-all neural network.** (A) For a 2-input comparator, scanning the $k_{cat}$ parameter values of the two proteases reveal that the circuit can correctly classify relative input levels as long as the protease corresponding to the larger input has a $k_{cat}$ value of at least 0.04/s. (B) For a 2-input comparator, bigger differences between the basal degradation rate (y-axis) and degron-based degradation rates (x-axis) result in more accurate classification (colorbar). (C) Deterministic simulation of the 2-input comparator with varying input levels. The circuit takes longer to classify inputs that are increasingly similar to each other, and can distinguish inputs within 10% of each other under reasonable time. (D) A fully connected 2-input neural network generates similar simulated dynamics as the comparator in both deterministic (E) and stochastic (F) simulations.

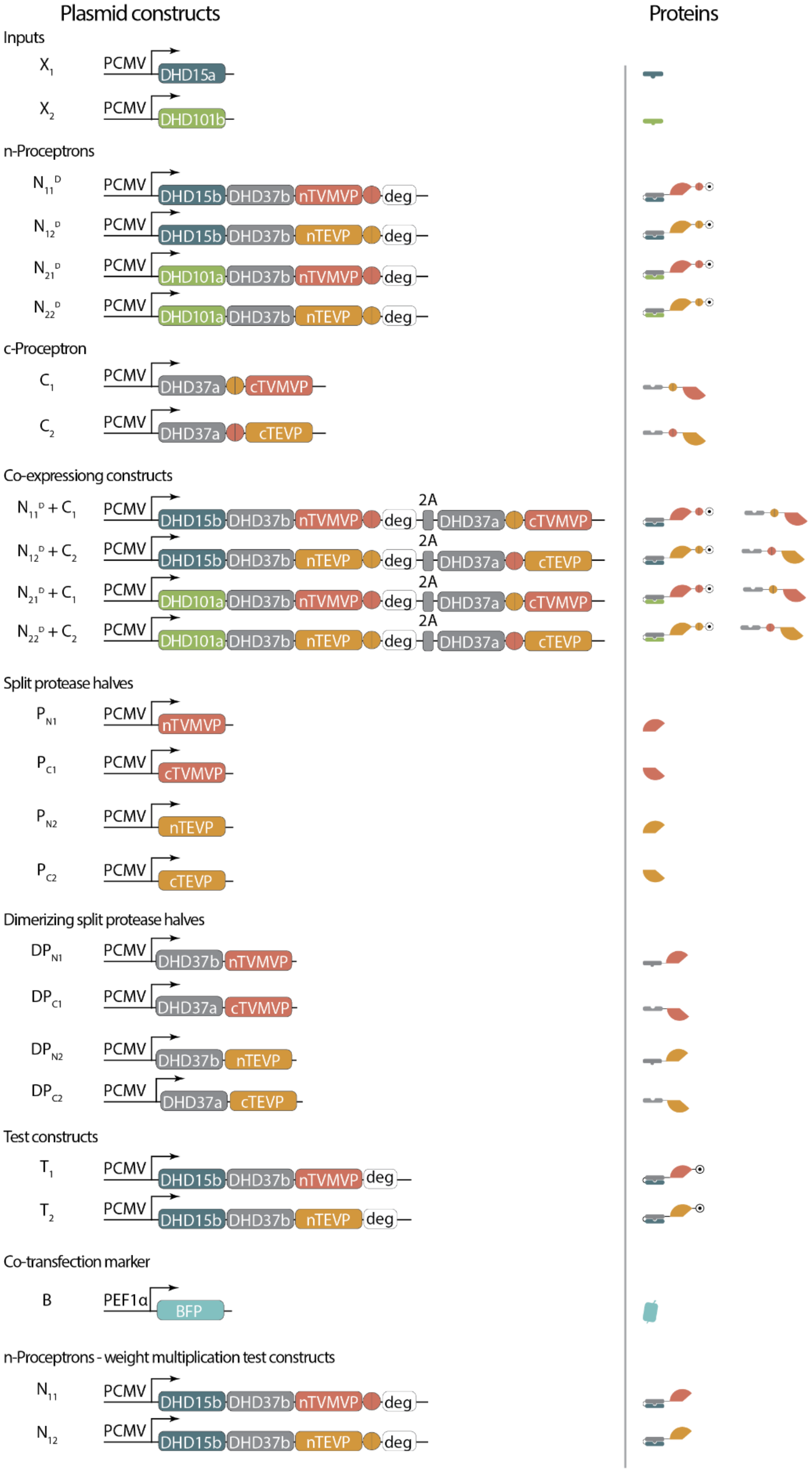

Plasmid constructs
Proteins
Inputs
$X_1$ PCMV DHD15a
$X_2$ PCMV DHD101b
n-Proceptrons
$N_{11}^D$ PCMV DHD15b DHD37b nTVMVP deg
$N_{12}^D$ PCMV DHD15b DHD37b nTEVP deg
$N_{21}^D$ PCMV DHD101a DHD37b nTVMVP deg
$N_{22}^D$ PCMV DHD101a DHD37b nTEVP deg
c-Proceptron
$C_1$ PCMV DHD37a cTVMVP
$C_2$ PCMV DHD37a cTEVP
Co-expressiong constructs
$N_{11}^D + C_1$ PCMV DHD15b DHD37b nTVMVP deg 2A DHD37a cTVMVP
$N_{12}^D + C_2$ PCMV DHD15b DHD37b nTEVP deg 2A DHD37a cTEVP
$N_{21}^D + C_1$ PCMV DHD101a DHD37b nTVMVP deg 2A DHD37a cTVMVP
$N_{22}^D + C_2$ PCMV DHD101a DHD37b nTEVP deg 2A DHD37a cTEVP
Split protease halves
$P_{N1}$ PCMV nTVMVP
$P_{C1}$ PCMV cTVMVP
$P_{N2}$ PCMV nTEVP
$P_{C2}$ PCMV cTEVP
Dimerizing split protease halves
$DP_{N1}$ PCMV DHD37b nTVMVP
$DP_{C1}$ PCMV DHD37a cTVMVP
$DP_{N2}$ PCMV DHD37b nTEVP
$DP_{C2}$ PCMV DHD37a cTEVP
Test constructs
$T_1$ PCMV DHD15b DHD37b nTVMVP deg
$T_2$ PCMV DHD15b DHD37b nTEVP deg
Co-transfection marker
B PEF1α BFP
n-Proceptrons - weight multiplication test constructs
$N_{11}$ PCMV DHD15b DHD37b nTVMVP deg
$N_{12}$ PCMV DHD15b DHD37b nTEVP deg

**Figure S2. Complete list of plasmids and the encoded protein constructs.** PCMV, human cytomegalovirus promoter. PEF1α, human elongation factor-1 α promoter. PGK, 3-phosphoglycerate kinase promoter. Deg, DHFR degron. BFP, blue fluorescent protein. Schematics of the resulting constructs are shown on the right.

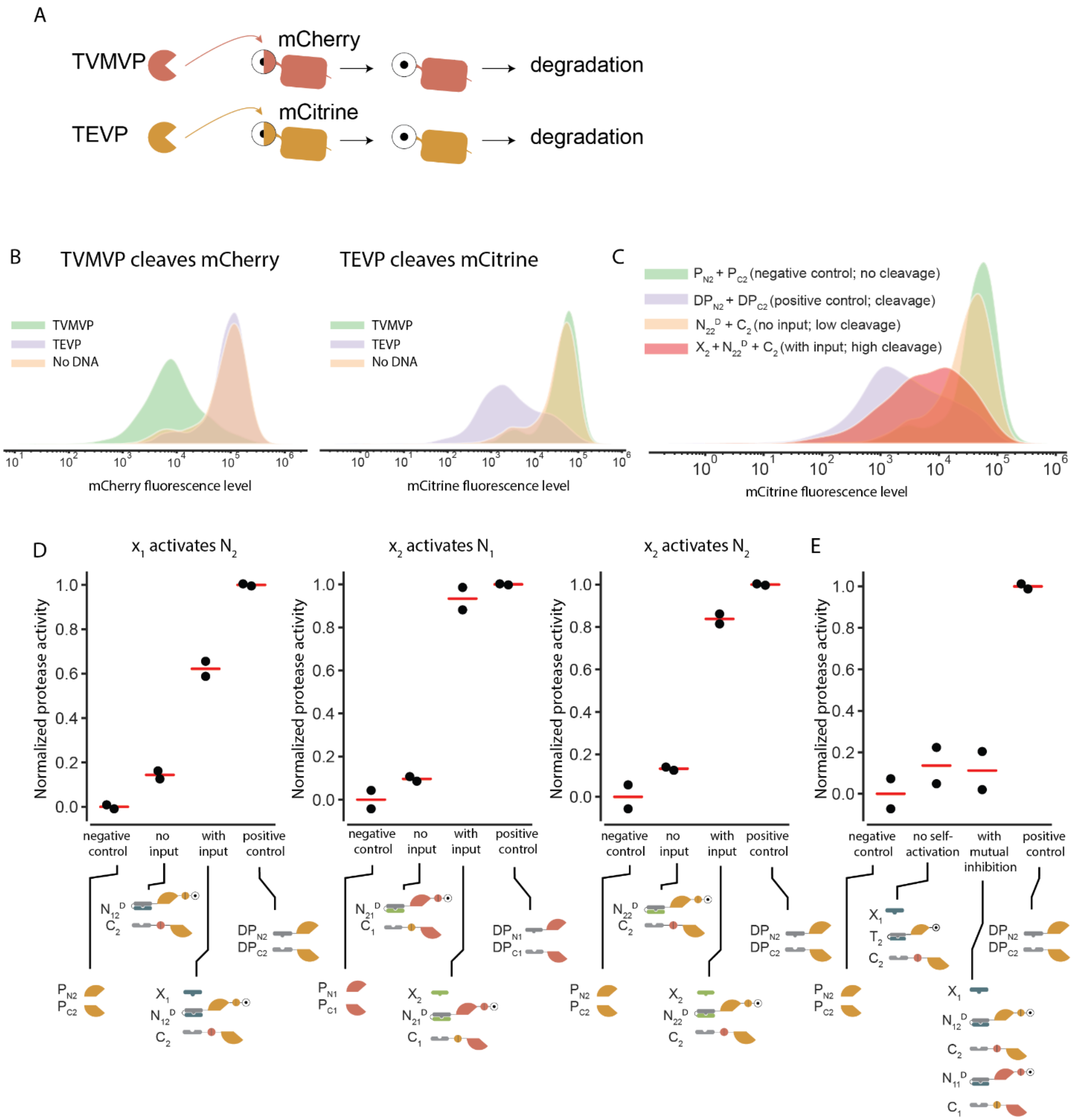

A
TVMVP
TEVP
mCherry
mCitrine
degradation
B
TVMVP cleaves mCherry
TEVP cleaves mCitrine
TVMVP
TEVP
No DNA
mCherry fluorescence level
mCitrine fluorescence level
C
P_N2 + P_C2 (negative control; no cleavage)
DP_N2 + DP_C2 (positive control; cleavage)
N_22^D + C_2 (no input; low cleavage)
X_2 + N_22^D + C_2 (with input; high cleavage)
mCitrine fluorescence level
D
x_1 activates N_2
x_2 activates N_1
x_2 activates N_2
E
Normalized protease activity
negative control
no input
with input
positive control
no self-activation
with mutual inhibition

**Figure S3. Experimental validation of the winner-take-all neural network.** (A) To read out protease activities, the reporter cell line (Figure 4B) expresses mCherry and mCitrine fluorescent proteins with N-end degrons that can be revealed upon protease cleavage by TVMVP and TEVP, respectively. Schematic shows how cleavage can expose degron (half-circle), to cause target protein degradation. (B) Flow cytometry histograms reveal that TVMVP and TEVP exclusively destabilize mCherry and mCitrine, respectively. (C) A representative flow cytometry histogram showing the activation of Node 2 by the $N_{22}D$ protein and its input $X_2$. (D) Activation of other nodes by their corresponding input proteins. (E) The node protein $N_{12}D$ undergoes self-activation, and its activities can be inhibited by the opposing node protein $N_{11}D$.

Table S1. Reaction rates used in simulations.

| Rates | Description | Value | Reference |
|---|---|---|---|
| $kon_1$ | on rate for the first step of cooperative DHD binding | $10^5$ $s^{-1}$ $M^{-1}$ | (*26*) |
| $koff_1$ | off rate for the first step of cooperative DHD binding | 100 $s^{-1}$ | (*26*) |
| $kon_2$ | on rate for the second step of cooperative DHD binding | $10^5$ $s^{-1}$ $M^{-1}$ | (*26*) |
| $koff_2$ | off rate for the second step of cooperative DHD binding | $10^{-4}$ $s^{-1}$ | (*26*) |
| $kon_p$ | on rate between proteases and substrates | $10^5$ $s^{-1}$ $M^{-1}$ | estimated |
| $koff_p$ | off rate between proteases and substrates | $10^{-4}$ $s^{-1}$ | estimated |
| $deg_{reg}$ | regular protein degradation rate | $10^{-5}$ $s^{-1}$ | (*39*) |
| $deg_{DHFR}$ | degradation rate for DHFR-tagged proteins | $10^{-3}$ $s^{-1}$ - $10^{-2}$ $s^{-1}$ | (*40*) |
| $k_{cat}$ | protease turnover number | 0.16 $s^{-1}$ | (*41*) |
| $V_{cell}$ | mammalian cell volume | $4*10^{-15}$ L | (*42*) |
| $k_{syn}$ | protein synthesis rate | 0.1-10 $s^{-1}$ | estimated |

Table S2. Proteins used in this study.

| Name used in modeling and simulation | Protein domains ("-" denotes a flexible linker) | Symbol | Sequence |
|---|---|---|---|
| $X_1$ | DHD15A | 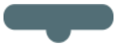 | TREELLRENIELAKE HIEIMREILELLQKM EELLEKARGADEDV AKTIKELLRRLKEIIE RNQRIAKEHEYIARE RSS |

| | | | |
|---|---|---|---|
| $X_2$ | DHD101B | 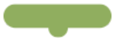 | GSDAYDLDRIVKEH<br>RRLVEEQRELVEEL<br>EKLVRRQEDHRVDK<br>KESHEILERLERIIRR<br>STRILTELEKLTDEF<br>ERRTR |
| $N_{11}^{D}$ | DHD15B-DHD37B-n<br>TVMVP-TVMVcs-D<br>HFR | 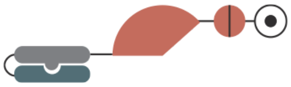 | GTERKLLERSRRLQ<br>EESKRLLDEMAEIM<br>RRIKKLLKKARGAD<br>EKVLDELRKIIERIRE<br>LLDRSRKIHERSEEI<br>AYKEEGSEGSGSE<br>GSGSDDKELDKLLD<br>TLEKILQTATKIIDDA<br>NKLLEKLRRSERKD<br>PKVVETYVELLKRH<br>EKAVKELLEIAKTHA<br>KKVEGSEGSGSEG<br>SSKALLKGVRDFNPI<br>SACVCLLENSSDGH<br>SERLFGIGFGPYIIA<br>NQHLFRRNNGELTI<br>KTMHGEFKVKNST<br>QLQMKPVEGRDIIVI<br>KMAKDFPPFPQKLK<br>FRQPTIKDRVCMVS<br>TNFQQSGGGSSET<br>VRFQSGSGSISLIAA<br>LAVDYVIGMENAMP<br>WNLPADLAWFKRN<br>TLNKPVIMGRHTWE<br>SIGRPLPGRKNIILS<br>SQPSTDDRVTWVK<br>SVDEAIAACGDVPEI<br>MVIGGGRVIEQFLP<br>KAQKLYLTHIDAEVE<br>GDTHFPDYEPDDW<br>ESVFSEFHDADAQN<br>SHSYCFEILERR |
| $N_{12}^{D}$ | DHD15B-DHD37B-n<br>TEVP-TEVcs-DHF<br>R | 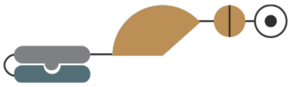 | GTERKLLERSRRLQ<br>EESKRLLDEMAEIM<br>RRIKKLLKKARGAD<br>EKVLDELRKIIERIRE<br>LLDRSRKIHERSEEI<br>AYKEEGSEGSGSE<br>GSGSDDKELDKLLD<br>TLEKILQTATKIIDDA<br>NKLLEKLRRSERKD |

| | | | |
|---|---|---|---|
| | | | PKVVETYVELLKRH<br>EKAVKELLEIAKTHA<br>KKVEGSEGSGSEG<br>SGESLFKGPRDYNP<br>ISSTICHLTNESDGH<br>TTSLYGIGFGPFIITN<br>KHLFRRNNGTLLVQ<br>SLHGVFKVKNTTTL<br>QQHLIDGRDMIIIRM<br>PKDFPPFPQKLKFR<br>EPQREERICLVTTNF<br>QTGGGSSENLYFQ<br>SGSGSISLIAALAVD<br>YVIGMENAMPWNL<br>PADLAWFKRNTLNK<br>PVIMGRHTWESIGR<br>PLPGRKNIILSSQPS<br>TDDRVTWVKSVDE<br>AIAACGDVPEIMVIG<br>GGRVIEQFLPKAQK<br>LYLTHIDAEVEGDTH<br>FPDYEPDDWESVF<br>SEFHDADAQNSHS<br>YCFEILERR |
| $N_{21}^{D}$ | DHD101A-DHD37B-nTVMVP-TVMVcs-DHFR | 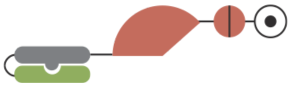 | DEKDYHRRLIEHLE<br>DLVRRHEELIKRQK<br>KVVEELERRGLDER<br>LRRVVDRFRRSSER<br>WEEVIERFRQVVDK<br>LRKSVEGSEGSGSE<br>GSGSDDKELDKLLD<br>TLEKILQTATKIIDDA<br>NKLLEKLRRSERKD<br>PKVVETYVELLKRH<br>EKAVKELLEIAKTHA<br>KKVEGSEGSGSEG<br>SSKALLKGVRDFNPI<br>SACVCLLENSSDGH<br>SERLFGIGFGPYIIA<br>NQHLFRRNNGELTI<br>KTMHGEFKVKNST<br>QLQMKPVEGRDIIVI<br>KMAKDFPPFPQKLK<br>FRQPTIKDRVCMVS<br>TNFQQSGGGSSET<br>VRFQSGSGSISLIAA<br>LAVDYVIGMENAMP<br>WNLPADLAWFKRN<br>TLNKPVIMGRHTWE |

| | | | |
|---|---|---|---|
| | | | SIGRPLPGRKNIILS<br>SQPSTDDRVTWVK<br>SVDEAIAACGDVPEI<br>MVIGGGRVIEQFLP<br>KAQKLYLTHIDAEVE<br>GDTHFPDYEPDDW<br>ESVFSEFHDADAQN<br>SHSYCFEILERR |
| $N_{22}^{D}$ | DHD101A-DHD37B-<br>nTEVP-TEVcs-DHF<br>R | 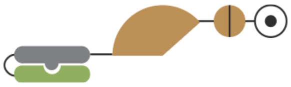 | DEKDYHRRLIEHLE<br>DLVRRHEELIKRQK<br>KVVEELERRGLDER<br>LRRVVDRFRRSSER<br>WEEVIERFRQVVDK<br>LRKSVEGSEGSGSE<br>GSGSDDKELDKLLD<br>TLEKILQTATKIIDDA<br>NKLLEKLRRSERKD<br>PKVVETYVELLKRH<br>EKAVKELLEIAKTHA<br>KKVEGSEGSGSEG<br>SGESLFKGPRDYNP<br>ISSTICHLTNESDGH<br>TTSLYGIGFGPFIITN<br>KHLFRRNNGTLLVQ<br>SLHGVFKVKNTTTL<br>QQHLIDGRDMIIIRM<br>PKDFPPFPQKLKFR<br>EPQREERICLVTTNF<br>QTGGGSSENLYFQ<br>SGSGSISLIAALAVD<br>YVIGMENAMPWNL<br>PADLAWFKRNTLNK<br>PVIMGRHTWESIGR<br>PLPGRKNIILSSQPS<br>TDDRVTWVKSVDE<br>AIAACGDVPEIMVIG<br>GGRVIEQFLPKAQK<br>LYLTHIDAEVEGDTH<br>FPDYEPDDWESVF<br>SEFHDADAQNSHS<br>YCFEILERR |
| $N_{11}$ | DHD15B-DHD37B-n<br>TVMVP | 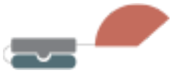 | GTERKLLERSRRLQ<br>EESKRLLDEMAEIM<br>RRIKKLLKKARGAD<br>EKVLDELRKIIERIRE<br>LLDRSRKIHERSEEI<br>AYKEEGSEGSGSE<br>GSGSDDKELDKLLD |

| | | | |
|---|---|---|---|
| | | | TLEKILQTATKIIDDA<br>NKLLEKLRRSERKD<br>PKVVETYVELLKRH<br>EKAVKELLEIAKTHA<br>KKVEGSEGSGSEG<br>SSKALLKGVRDFNPI<br>SACVCLLENSSDGH<br>SERLFGIGFGPYIIA<br>NQHLFRRNNGELTI<br>KTMHGEFKVKNST<br>QLQMKPVEGRDIIVI<br>KMAKDFPPFPQKLK<br>FRQPTIKDRVCMVS<br>TNFQQS |
| $N_{12}$ | DHD15B-DHD37B-n<br>TEVP | 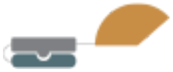 | GTERKLLERSRRLQ<br>EESKRLLDEMAEIM<br>RRIKKLLKKARGAD<br>EKVLDELRKIIERIRE<br>LLDRSRKIHERSEEI<br>AYKEEGSEGSGSE<br>GSGSDDKELDKLLD<br>TLEKILQTATKIIDDA<br>NKLLEKLRRSERKD<br>PKVVETYVELLKRH<br>EKAVKELLEIAKTHA<br>KKVEGSEGSGSEG<br>SGESLFKGPRDYNP<br>ISSTICHLTNESDGH<br>TTSLYGIGFGPFIITN<br>KHLFRRNNGTLLVQ<br>SLHGVFKVKNTTTL<br>QQHLIDGRDMIIIRM<br>PKDFPPFPQKLKFR<br>EPQREERICLVTTNF<br>QT |
| $C_1$ | DHD37A-TEVcs-cT<br>VMVP | 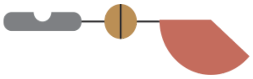 | DSDEHLYKLKTFLE<br>NLRRHLDRLDKHIK<br>QLRDILSENPEDER<br>VKDVIDLSERSVRIV<br>KTVIKIFEDSVRKKE<br>GSEGSGSEGSENL<br>YFQSGSKSVSSLVS<br>ESSHIVHKEDTSFW<br>QHWITTKDGQCGS<br>PLVSIIDGNILGIHSL<br>THTTNGSNYFVEFP<br>EKFVATYLDAADGW<br>CKNWKFNADKISW |

| | | | |
|---|---|---|---|
| | | | GSFTLVEDAPEDDF<br>MAKKTVAAIMD |
| $C_2$ | DHD37A-TVMVcs-c<br>TEVP | 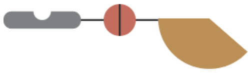 | DSDEHLYKLKTFLE<br>NLRRHLDRLDKHIK<br>QLRDILSENPEDER<br>VKDVIDLSERSVRIV<br>KTVIKIFEDSVRKKE<br>GSEGSGSEGSETV<br>RFQSGSKSMSSMV<br>SDTSCTFPSSDGIF<br>WKHWIQTKDGQCG<br>SPLVSTRDGFIVGIH<br>SASNFTNTNNYFTS<br>VPKNFMELLTNQEA<br>QQWVSGWRLNADS<br>VLWGGHKVFMVKP<br>EEPFQPVKEATQLM<br>NS |
| $P_{N1}$ | nTVMVP | 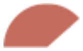 | SKALLKGVRDFNPIS<br>ACVCLLENSSDGHS<br>ERLFGIGFGPYIIAN<br>QHLFRRNNGELTIK<br>TMHGEFKVKNSTQL<br>QMKPVEGRDIIVIKM<br>AKDFPPFPQKLKFR<br>QPTIKDRVCMVSTN<br>FQQS |
| $P_{C1}$ | cTVMVP | 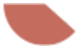 | KSVSSLVSESSHIVH<br>KEDTSFWQHWITTK<br>DGQCGSPLVSIIDG<br>NILGIHSLTHTTNGS<br>NYFVEFPEKFVATYL<br>DAADGWCKNWKFN<br>ADKISWGSFTLVED<br>APEDDFMAKKTVAA<br>IMD |
| $P_{N2}$ | nTEVP | 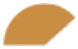 | GESLFKGPRDYNPI<br>SSTICHLTNESDGH<br>TTSLYGIGFGPFIITN<br>KHLFRRNNGTLLVQ<br>SLHGVFKVKNTTTL<br>QQHLIDGRDMIIIRM<br>PKDFPPFPQKLKFR<br>EPQREERICLVTTNF<br>QT |

| | | | |
|---|---|---|---|
| $P_{C2}$ | cTEVP | 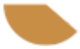 | KSMSSMVSDTSCTFPSSDGIFWKHWIQTKDGQCGSPLVSTRDGFIVGIHSASNFTNTNNYFTSVPKNFMELLTNQEAQQWVSGWRLNADSVLWGGHKVFMVKPEEPFQPVKEATQLMN |
| $DP_{N1}$ | DHD37B-nTVMVP | 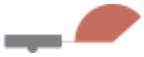 | GSDDKELDKLLDTLEKILQTATKIIDDANKLLEKLRRSERKDPKVVETYVELLKRHEKAVKELLEIAKTHAKKVEGSEGSGSEGSSKALLKGVRDFNPISACVCLLENSSDGHSERLFGIGFGPYIIANQHLFRRNNGELTIKTMHGEFKVKNSTQLQMKPVEGRDIIVIKMAKDFPPFPQKLKFRQPTIKDRVCMVSTNFQQS |
| $DP_{C1}$ | DHD37A-cTVMVP | 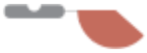 | DSDEHLYKLKTFLENLRRHLDRLDKHIKQLRDILSENPEDERVKDVIDLSERSVRIVKTVIKIFEDSVRKKEGSEGSGSEGSKSVSSLVSESSHIVHKEDTSFWQHWITTKDGQCGSPLVSIIDGNILGIHSLTHTTNGSNYFVEFPEKFVATYLDAADGWCKNWKFNADKISWGSFTLVEDAPEDDFMAKKTVAAIMD |
| $DP_{N2}$ | DHD37B-nTEVP | 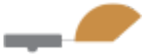 | GSDDKELDKLLDTLEKILQTATKIIDDANKLLEKLRRSERKDPKVVETYVELLKRHEKAVKELLEIAKTHAKKVEGSEGSGSEGSGESLFKGPRDYNPISSTICHLTNESDGHTT |

| | | | |
|---|---|---|---|
| | | | SLYGIGFGPFIITNKHLFRRNNGTLLVQSLHGVFKVKNTTTLQQHLIDGRDMIIIRMPKDFPPFPQKLKFREPQREERICLVTTNFQT |
| $DP_{C2}$ | DHD37A-cTEVP | 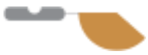 | DSDEHLYKLKTFLENLRRHLDRLDKHIKQLRDILSENPEDERVKDVIDLSERSVRIVKTVIKIFEDSVRKKEGSEGSGSEGSKSMSSMVSDTSCTFPSSDGIFWKHWIQTKDGQCGSPLVSTRDGFIVGIHSASNFTNTNNYFTSVPKNFMELLTNQEAQQWVSGWRLNADSVLWGGHKVFMVKPEEPFQPVKEATQLMN |
| $T_1$ | DHD15B-DHD37B-nTVMVP-DHFR | 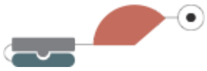 | GTERKLLERSRRLQEESKRLLDEMAEIMRRIKKLLKKARGADEKVLDELRKIIERIRELLDRSRKIHERSEEIAYKEEGSEGSGSEGSGSDDKELDKLLDTLEKILQTATKIIDDANKLLEKLRRSERKDPKVVETYVELLKRHEKAVKELLEIAKTHAKKVEGSEGSGSEGSSKALLKGVRDFNPISACVCLLENSSDGHSERLFGIGFGPYIIANQHLFRRNNGELTIKTMHGEFKVKNSTQLQMKPVEGRDIIVIKMAKDFPPFPQKLKFRQPTIKDRVCMVSTNFQQSGSGSISLIAALAVDYVIGMENAMPWNLPADLAWFKRNTLNKPVIMGRHTWESIGRPLPGRKNIILSSQPSTDDRVTWV |

| | | | |
|---|---|---|---|
| | | | KSVDEAIAACGDVP<br>EIMVIGGGRVIEQFL<br>PKAQKLYLTHIDAEV<br>EGDTHFPDYEPDD<br>WESVFSEFHDADA<br>QNSHSYCFEILERR |
| $T_2$ | DHD15B-DHD37B-nTEVP-DHFR | 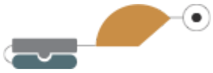 | GTERKLLERSRRLQ<br>EESKRLLDEMAEIM<br>RRIKKLLKKARGAD<br>EKVLDELRKIIERIRE<br>LLDRSRKIHERSEEI<br>AYKEEGSEGSGSE<br>GSGSDDKELDKLLD<br>TLEKILQTATKIIDDA<br>NKLLEKLRRSERKD<br>PKVVETYVELLKRH<br>EKAVKELLEIAKTHA<br>KKVEGSEGSGSEG<br>SGESLFKGPRDYNP<br>ISSTICHLTNESDGH<br>TTSLYGIGFGPFIITN<br>KHLFRRNNGTLLVQ<br>SLHGVFKVKNTTTL<br>QQHLIDGRDMIIIRM<br>PKDFPPFPQKLKFR<br>EPQREERICLVTTNF<br>QTGGGSSISLIAALA<br>VDYVIGMENAMPW<br>NLPADLAWFKRNTL<br>NKPVIMGRHTWESI<br>GRPLPGRKNIILSSQ<br>PSTDDRVTWVKSV<br>DEAIAACGDVPEIM<br>VIGGGRVIEQFLPKA<br>QKLYLTHIDAEVEGD<br>THFPDYEPDDWES<br>VFSEFHDADAQNSH<br>SYCFEILERR |
| N/A | BFP | 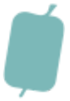 | VSKGEELIKENMHM<br>KLYMEGTVDNHHFK<br>CTSEGEGKPYEGT<br>QTMRIKVVEGGPLP<br>FAFDILATSFLYGSK<br>TFINHTQGIPDFFKQ<br>SFPEGFTWERVTTY<br>EDGGVLTATQDTSL<br>QDGCLIYNVKIRGV<br>NFTSNGPVMQKKTL |

| | | | |
|---|---|---|---|
| | | | GWEAFTETLYPADG<br>GLEGRNDMALKLVG<br>GSHLIANAKTTYRS<br>KKPAKNLKMPGVYY<br>VDYRLERIKEANNE<br>TYVEQHEVAVARYC<br>DLPSKLGHKLN |
| N/A | mCherry | 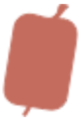 | GGVSKGEEDNMAII<br>KEFMRFKVHMEGS<br>VNGHEFEIEGEGEG<br>RPYEGTQTAKLKVT<br>KGGPLPFAWDILSP<br>QFMYGSKAYVKHPA<br>DIPDYLKLSFPEGFK<br>WERVMNFEDGGVV<br>TVTQDSSLQDGEFI<br>YKVKLRGTNFPSDG<br>PVMQKKTMGWEAS<br>SERMYPEDGALKG<br>EIKQRLKLKDGGHY<br>DAEVKTTYKAKKPV<br>QLPGAYNVNIKLDIT<br>SHNEDYTIVEQYER<br>AEGRHSTGGMDEL<br>YKS |
| N/A | mCitrine | 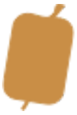 | VSKGEELFTGVVPIL<br>VELDGDVNGHKFSV<br>SGEGEGDATYGKLT<br>LKFICTTGKLPVPW<br>PTLVTTFGYGLMCF<br>ARYPDHMKQHDFF<br>KSAMPEGYVQERTI<br>FFKDDGNYKTRAEV<br>KFEGDTLVNRIELKG<br>IDFKEDGNILGHKLE<br>YNYNSHNVYIMADK<br>QKNGIKVNFKIRHNI<br>EDGSVQLADHYQQ<br>NTPIGDGPVLLPDN<br>HYLSYQSALSKDPN<br>EKRDHMVLLEFVTA<br>AGITLGMDELYKS |
| N/A | TVMVcs-Degron-mCherry | 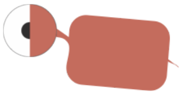 | ETVRFQYHKSGAW<br>KLPVSLVKGGTSVS<br>KGEEDNMAIIKEFM<br>RFKVHMEGSVNGH<br>EFEIEGEGEGRPYE |

| | | | |
|---|---|---|---|
| | | | GTQTAKLKVTKGGPLPFAWDILSPQFMYGSKAYVKHPADIPDYLKLSFPEGFKWERVMNFEDGGVVTVTQDSSLQDGEFIYKVKLRGTNFPSDGPVMQKKTMGWEASSERMYPEDGALKGEIKQRLKLKDGGHYDAEVKTTYKAKKPVQLPGAYNVNIKLDITSHNEDYTIVEQYERAEGRHSTGGMDELYKS |
| N/A | TEVcs-Degron-mCitrine | 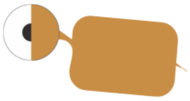 | ENLYFQYHKSGAWKLPVSLVKGGGSVSKGEELFTGVVPILVELDGDVNGHKFSVSGEGEGDATYGKLTLKFICTTGKLPVPWPTLVTTFGYGLMCFARYPDHMKQHDFFKSAMPEGYVQERTIFFKDDGNYKTRAEVKFEGDTLVNRIELKGIDFKEDGNILGHKLEYNYNSHNVYIMADKQKNGIKVNFKIRHNIEDGSVQLADHYQQNTPIGDGPVLLPDNHYLSYQSALSKDPNEKRDHMVLLEFVTAAGITLGMDELYKS |

Table S3. Plasmids and mRNAs used in transfection experiments.

| Figure panel | Plasmids/mRNAs used |
|---|---|
| 3C | 1. $DP_{N1}$, 50 ng; $DP_{C1}$, 50 ng<br>2. $N_{11}^{D}$, 50 ng; $C_1$ 50 ng<br>3. $N_{11}^{D}$, 50 ng; $C_1$ 50 ng; $X_1$, 400ng<br>4. $X_1$, 50 ng; $T_1$, 50 ng; $C_1$, 50 ng<br>5. $X_1$, 50 ng; ($N_{11}^{D}$+ $C_1$), 50 ng; ($N_{12}^{D}$ + $C_2$), 450 ng |

<table>
<tr><td></td><td>6. $DP_{N1}$, 50 ng; $DP_{C1}$, 50 ng</td></tr>
<tr><td>3D</td><td>mRNAs<br>$X_1$, 200 ng; $DP_{C1}$ 200 ng; $DP_{C2}$ 200 ng<br><br>$N_{11}$ and $N_{12}$ are varied base on the following table:<br><br>
<table>
<tr><td>$N_{11}$ (ng)</td><td>$N_{12}$ (ng)</td></tr>
<tr><td>500</td><td>0</td></tr>
<tr><td>450</td><td>50</td></tr>
<tr><td>400</td><td>100</td></tr>
<tr><td>300</td><td>200</td></tr>
<tr><td>250</td><td>250</td></tr>
<tr><td>200</td><td>300</td></tr>
<tr><td>100</td><td>400</td></tr>
<tr><td>50</td><td>450</td></tr>
<tr><td>0</td><td>500</td></tr>
</table>
</td></tr>
<tr><td>3E</td><td>($N_{11}^D$+ $C_1$), 50 ng; ($N_{21}^D$+ $C_1$), 3.3 ng;<br>($N_{12}^D$+ $C_2$), 5 ng; ($N_{22}^D$+ $C_2$), 33 ng;<br><br>Amount of input proteins are indicated in the figure</td></tr>
<tr><td>3F</td><td>($N_{11}^D$+ $C_1$), 50 ng; ($N_{21}^D$+ $C_1$), 50 ng;<br>($N_{12}^D$+ $C_2$), 16 ng; ($N_{22}^D$+ $C_2$), 16 ng;<br><br>Amount of input proteins are indicated in the figure</td></tr>
<tr><td>3G</td><td>5:1 -- ($N_{11}^D$+ $C_1$), 50 ng; ($N_{22}^D$+ $C_2$), 10 ng<br>2:1 -- ($N_{11}^D$+ $C_1$), 50 ng; ($N_{22}^D$+ $C_2$), 25 ng<br>1.5:1 -- ($N_{11}^D$+ $C_1$), 50 ng; ($N_{22}^D$+ $C_2$), 33 ng<br>1:1 -- ($N_{11}^D$+ $C_1$), 50 ng; ($N_{22}^D$+ $C_2$), 50 ng<br>1:1.5 -- ($N_{11}^D$+ $C_1$), 33 ng; ($N_{22}^D$+ $C_2$), 50 ng<br>1:2 -- ($N_{11}^D$+ $C_1$), 25 ng; ($N_{22}^D$+ $C_2$), 50 ng<br>1:5 -- ($N_{11}^D$+ $C_1$), 10 ng; ($N_{22}^D$+ $C_2$), 50 ng</td></tr>
</table>

| | |
|---|---|
| | Amount of input proteins are indicated in the figure |
| S4B | 7. $DP_{N1}$, 50 ng; $DP_{C1}$, 50 ng<br>8. $DP_{N2}$, 50 ng; $DP_{C2}$, 50 ng |
| S4C | 1. $P_{N2}$, 50 ng; $P_{C2}$, 50 ng<br>2. $N_{22}^{D}$, 50 ng; $C_2$ 50 ng<br>3. $N_{22}^{D}$, 50 ng; $C_2$ 50 ng; $X_2$, 400ng<br>4. $DP_{N2}$, 50 ng; $DP_{C2}$, 50 ng |
| S4D | Left:<br>1. $P_{N2}$, 50 ng; $P_{C2}$, 50 ng<br>2. $N_{12}^{D}$, 50 ng; $C_2$ 50 ng<br>3. $N_{12}^{D}$, 50 ng; $C_2$ 50 ng; $X_1$, 400ng<br>4. $DP_{N2}$, 50 ng; $DP_{C2}$, 50 ng<br><br>Middle:<br>9. $P_{N1}$, 50 ng; $P_{C1}$, 50 ng<br>10. $N_{21}^{D}$, 50 ng; $C_1$ 50 ng<br>11. $N_{21}^{D}$, 50 ng; $C_1$ 50 ng; $X_2$, 400ng<br>12. $DP_{N1}$, 50 ng; $DP_{C1}$, 50 ng<br><br>Right:<br>5. $P_{N2}$, 50 ng; $P_{C2}$, 50 ng<br>6. $N_{22}^{D}$, 50 ng; $C_2$ 50 ng<br>7. $N_{22}^{D}$, 50 ng; $C_2$ 50 ng; $X_2$, 400ng<br>8. $DP_{N2}$, 50 ng; $DP_{C2}$, 50 ng |
| S4E | 1. $P_{N2}$, 50 ng; $P_{C2}$, 50 ng<br>2. $X_1$, 50 ng; $T_2$, 50 ng; $C_2$, 50 ng<br>3. $X_1$, 50 ng; ($N_{12}^{D}$ + $C_2$), 50 ng; ($N_{11}^{D}$+ $C_1$), 450 ng<br>4. $DP_{N2}$, 50 ng; $DP_{C2}$, 50 ng |
